# Functional anatomy of the full length CXCR4-CXCL12 complex systematically dissected by quantitative model-guided mutagenesis

**DOI:** 10.1101/2020.01.21.913772

**Authors:** Bryan S. Stephens, Tony Ngo, Irina Kufareva, Tracy M. Handel

**Affiliations:** Skaggs School of Pharmacy and Pharmaceutical Sciences, University of California, San Diego, La Jolla 92093, CA

## Abstract

Due to their prominent role in development and infamy in cancer and HIV, the chemokine receptor CXCR4 and its ligand, CXCL12, have been the subject of numerous structural and functional studies. Nevertheless, a high resolution structure of the CXCR4-CXCL12 complex has not been reported. Even with several alternative computational models of the complex at hand, the relative contributions of different interaction epitopes to ligand binding, ligand selectivity and signaling are not readily apparent. Here, building upon our latest structural model, we employed a systematic mutagenesis strategy to dissect the functional anatomy of the of CXCR4-CXCL12 complex. Key charge swap mutagenesis experiments supported pairwise interactions between oppositely charged residues in the receptor and chemokine, confirming the accuracy of the predicted orientation of the chemokine relative to the receptor, while also providing insight into ligand selectivity. Progressive deletion of N-terminal residues revealed an unexpected contribution of the receptor N-terminus to chemokine signaling; this finding challenges a longstanding “two-site” hypothesis about the essential features of the receptor-chemokine interaction where the N-terminus is purported to only contribute to binding affinity. The results suggest that while the interaction of the chemokine N-terminus with the receptor binding pocket is the key driver of signaling, the signaling amplitude depends on the extent to which the receptor N-terminus binds the chemokine. Along with systematic characterization of other epitopes, the current data allow us to propose a comprehensive experimentally-consistent structural model for how the chemokine binds CXCR4 and initiates signal transmission through the receptor TM domain.

**One sentence summary:** A systematic structure-guided mutagenesis study of chemokine receptor CXCR4 reveals novel insights into epitopes regulating ligand recognition, ligand specificity and CXCL12-mediated signaling.

## Introduction

Chemokine receptors are members of the Class A family G protein-coupled receptors (GPCRs) that are best known for their role in controlling the migration of cells, particularly leukocytes in the context of immune system function. They are activated by chemokines, small 8-10 kDa secreted proteins, via a mechanism that has long been described as involving “two sites” or “two steps” (*1–5*). According to the two-site mechanism, the globular domain of the chemokine binds to the N-terminus of a receptor (referred to as chemokine recognition site 1, CRS1) and contributes primarily to the stability of the complex whereas the N-terminus of the chemokine binds in the transmembrane (TM) domain binding pocket of the receptor (chemokine recognition site 2, CRS2) to activate signaling (*6*). The distinction between these two sites arose from the general observation that mutations in chemokine N-termini produce a disproportionately large effect on receptor signaling efficacy compared to mutations in the chemokine globular domains (*7, 8*), with corresponding trends observed for chimeric rearrangements (*1*) or mutations (*9*) of the corresponding CRS2 and CRS1 regions of the receptors. In fact, single point mutations or modifications of chemokine N-termini can completely alter receptor pharmacology, producing antagonists and even superagonists in many cases (*2, 7, 10-13*).

In 2015, our group solved the structure of the human CXC chemokine receptor 4 (CXCR4), in complex with vMIP-II, a viral CC chemokine antagonist from human herpesvirus, HHV8 (*14*). The CXCR4-vMIP-II structure confirmed the presence of CRS1 and CRS2 interactions as expected from the two-site model, but also revealed an intermediate region, CRS1.5, that bridges CRS1 and CRS2 and contributes to a contiguous interaction interface between the chemokine and receptor. Structures of three other complexes have also been determined - those of the virally encoded receptor US28 in complex with the human chemokine CX3CL1 and variants (*15, 16*), and that of the human chemokine receptor CCR5 bound to [5P7]CCL5, the engineered antagonist variant of human CCL5 (*17*). All crystallized complexes feature a similar contiguous interaction interface involving CRS1, CRS1.5 and CRS2, suggesting that these epitopes constitute an interaction architecture that is conserved in the chemokine receptor family. The structures also suggest that CRS1.5 acts as a pivot point that allows the relative orientations of the chemokine and receptor to differ between complexes, thereby contributing to ligand recognition and signaling specificity (*17*).

Despite being one of the most intensely studied chemokine receptors, initially because of its role as a cofactor for HIV infection (*18–20*) and subsequently because of its widespread role in cancer (*21–23*), a structure of CXCR4 in complex with its endogenous chemokine ligand CXCL12 has not yet been determined. Several computational models (*24–29*), along with our own (*14, 30, 31*) have been put forward, but significant differences in the proposed models highlight the need for experimental validation and refinement. Additionally, experimental data are required to understand how the structure of the complex translates into receptor activation, which is poorly understood, even for this well-studied receptor. There are several likely reasons for this. Prior mutational studies, although valuable, have often been focused on limited sets of mutations and originated from different laboratories using different techniques. Moreover, the reports of mutation effects have often been based on single point assays rather than full concentration response curves, and both single point and concentration response experiments are typically carried out without accounting for changes in expression levels, which can significantly influence the results. Finally, the most frequently used readout of CXCR4 activation, intracellular calcium mobilization, is subject to signal amplification that can mask the effect of mutations.

Here, we functionally dissect the signaling role of various features proposed in our latest computational model of CXCR4-CXCL12. Our approach includes reciprocal charge reversal (“charge swap”) rescue-of-function mutants in addition to single point mutants to provide evidence for the proposed orientation of the chemokine relative to the receptor. In addition, the model proposes an interface between the full receptor N-terminus and chemokine, which is not resolved in any chemokine receptor crystal structures solved to date. Progressive deletion of the N-terminus caused diminished β-arrestin and G protein recruitment, which was surprising given that the N-terminus has been purported to be primarily an affinity determinant. Building on our prior studies (*14, 30–32*), the current data allows us to propose a comprehensive experimentally-consistent structural framework explaining how the chemokine binds CXCR4 and initiates signal transmission through the receptor TM domain. The data also add to accumulating evidence suggesting that receptor-chemokine interactions are more complex than implied by the two-site mechanism and that residues outside of CRS2 can play an important role in CXCR4 activation.

## Results

### Full length model of the CXCR4-CXCL12 signaling complex

A model of the complex between full-length CXCR4 and CXCL12 (Fig. 1A) was produced via an integrated approach that combines homology modeling and flexible molecular docking with experimentally derived restraints from disulfide crosslinking (*31*). The architecture of the complex is consistent with that of all three crystallized receptor-chemokine complexes (*14, 15, 17*). It features the CRS1 interaction where the N-terminal residues 21-sYDSMKE-26 of the receptor bind in the groove formed by the “N-loop” and “40s loop” of CXCL12, and the CRS2 interaction where the flexible N-terminus of CXCL12 (NH_3_^+^-1-KPVSLSYR-8) reaches into the TM domain pocket of CXCR4, making contacts with the critical residues from the so-called “engagement layer” (Asp^97^(2.63), Asp^187^(ECL2), and Asp^262^(6.58); Ballesteros and Weinstein numbering in parenthesis) and “signal initiation layer” (Trp^94^(2.60), Tyr^116^(3.32), and Glu^288^(7.39)) (*30*). The two epitopes are joined by the CRS1.5 region (*14*) where Pro^27^(NT) and Cys^28^(NT) of CXCR4 pack against the first disulfide (Cys^9^-Cys^50^) of CXCL12 and its β_3_-strand in a conserved manner that has been observed not only in all three crystallized receptor-chemokine complexes (*17*), but also across multiple chemokine-binding proteins that are unrelated to receptors (*33, 34*). This suggests that CRS1.5 is an anchor point for various proteins interacting with chemokines. The conformations and interactions in these epitopes originate from our previously published partial model that featured CXCR4 residues 21-304 (*14*), and have been refined in (*31*).

**Fig. 1.**
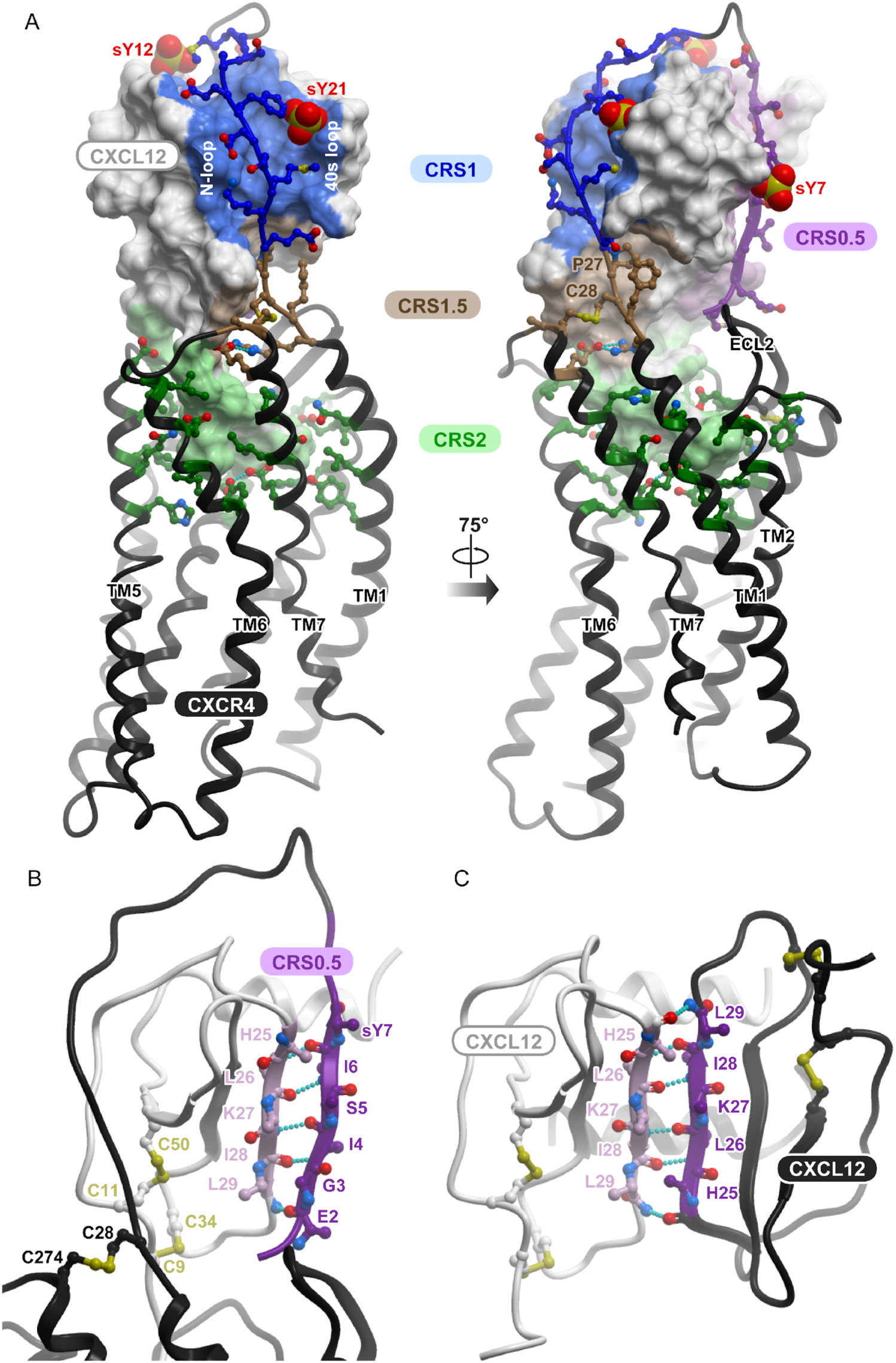
The computational model of the CXCR4-CXCL12 complex used to guide and interpret the experiments in this work. (**A**) The entire model is viewed along the plane of the membrane. The chemokine is shown as a surface mesh, the receptor as a black ribbon. The distinct interaction epitopes discussed in the paper are labeled. (**B**) The proposed CRS0.5 interaction involves an anti-parallel β-sheet between the distal N-terminus of the receptor and the β1 strand of the chemokine. (**C**) The proposed CRS0.5 interaction between the receptor and the chemokine closely mimics the interaction between CXCL12 monomers in the dimer (PDB 3gv3).

Novel to the present model is the complete N-terminus of CXCR4, including residues 1-20, a region that has not been resolved in any of the chemokine receptor-chemokine complex crystal structures (*14, 15, 17*). Prior mutagenesis studies have suggested that the entire N-terminus interacts with the chemokine (*35, Wescott, 2016 #2108*) and highlighted the roles of three sulfated tyrosines (sTyr^7^, sTyr^12^ and sTyr^21^) that affect binding affinity of the chemokine for the receptor (*36, 37*). The present model (Fig. 1A) suggests that the N-terminus of CXCR4 continues on from its position in the CXCR4 crystal structure and “wraps around” the chemokine, engaging residues from the chemokine 3_10_ helix and the C-terminal helix; it also suggests that the distal N-terminal residues 1-MEGISIsY-7 form an anti-parallel β-sheet with the β_1_-strand of the chemokine (Fig. 1B) in a manner that largely mimics the intermolecular packing in CXCL12 homodimers (Fig. 1C, *38*). These residues belong to an interaction epitope referred to as CRS0.5 (*39*).

Consistent with the abundance of charged residues in both CXCR4 and CXCL12, the intermolecular interaction in the model is mediated by numerous electrostatic contacts. Salt bridges between the N-terminal amine of the chemokine and CXCR4 Asp^97^(2.63), the side chain of CXCL12 Lys^1^ and CXCR4 Glu^288^(7.39), CXCL12 Arg^8^ and CXCR4 Asp^262^(6.58) form key electrostatic anchors in CRS2. The salt bridge between CXCR4 Glu^277^(7.28) and CXCL12 Arg^12^ (N-loop) supports the orientation of CXCL12 relative to CXCR4 in the CRS1.5 interaction epitope. These and other key oppositely charged residue pairs predicted by the model to be in contact are summarized in Table 1.

**Table 1.** Key predicted charge interactions in CRS1, 1.5, and 2 of the CXCR4-CXCL12 complex.

| **CXCR4** | **CXCL12** | **Epitope** |
| --- | --- | --- |
| Asp<sup>97</sup>(2.63) | NH<sub>3</sub><sup>+</sup> | CRS2 – engagement layer (30) |
| Glu<sup>288</sup>(7.39) | Lys<sup>1</sup> | CRS2 – initiation layer (30) |
| Asp<sup>262</sup>(6.58) | Arg<sup>8</sup> | CRS2 – engagement layer (30) |
| Glu<sup>277</sup>(7.28) | Arg<sup>12</sup> | CRS1.5 |
| Glu<sup>26</sup>(NT) | Arg<sup>47</sup> (40s loop) | CRS1 |
| Lys<sup>25</sup>(NT) | Glu<sup>15</sup> (N-loop) | CRS1 |
| Asp<sup>22</sup>(NT) | His<sup>17</sup> (N-loop) | CRS1 |
| sTyr<sup>21</sup>(NT) | Asn<sup>22</sup> (3<sub>10</sub> helix), Asn<sup>44</sup> and<br>Asn<sup>45</sup> (40s loop) | CRS1 |
| Asp<sup>20</sup>(NT) | Lys<sup>56</sup> | CRS1 |

Altogether, the model suggests that the interface between the receptor and the chemokine is compositionally complex. At a minimum, four constituent epitopes can be clearly identified: CRS0.5, CRS1, CRS1.5, and CRS2. Compared to the broadly defined roles of the best-known epitopes, CRS1 and CRS2, almost nothing is known about the role of CRS1.5 contacts in binding and signaling, and the newly proposed CRS0.5 epitope has never been studied. Moreover, the contributions of the individual residue contacts in these four epitopes (as well as other regions) to the affinity or pharmacology of the CXCR4-CXCL12 complex are unclear. Guided by the model, we set out to quantitatively dissect the anatomy of the CXCR4-CXCL12 interface, and the roles of its constituent epitopes in triggering downstream signaling.

### Mutagenesis strategy to quantitatively assess the signaling capacity of CXCR4 mutants

Prior mutagenesis studies probing CXCR4-CXCL12 interactions studied herein have almost exclusively focused on G protein signaling with little attention given to the involvement of β-arrestins, which are also important in CXCR4 function (*40–42*). Moreover, these studies often rely on single-CXCL12 concentration measurements, are sometimes hindered in interpretation by varying mutant expression levels, and almost always rely on second messenger signaling data that are subject to signal amplification (*43–45*). While valuable, these studies do not provide a consistent, uniform, and quantitative assessment of mutants, making it difficult to integrate them into a cohesive model of CXCR4 signaling (Table 2).

**Table 2.**
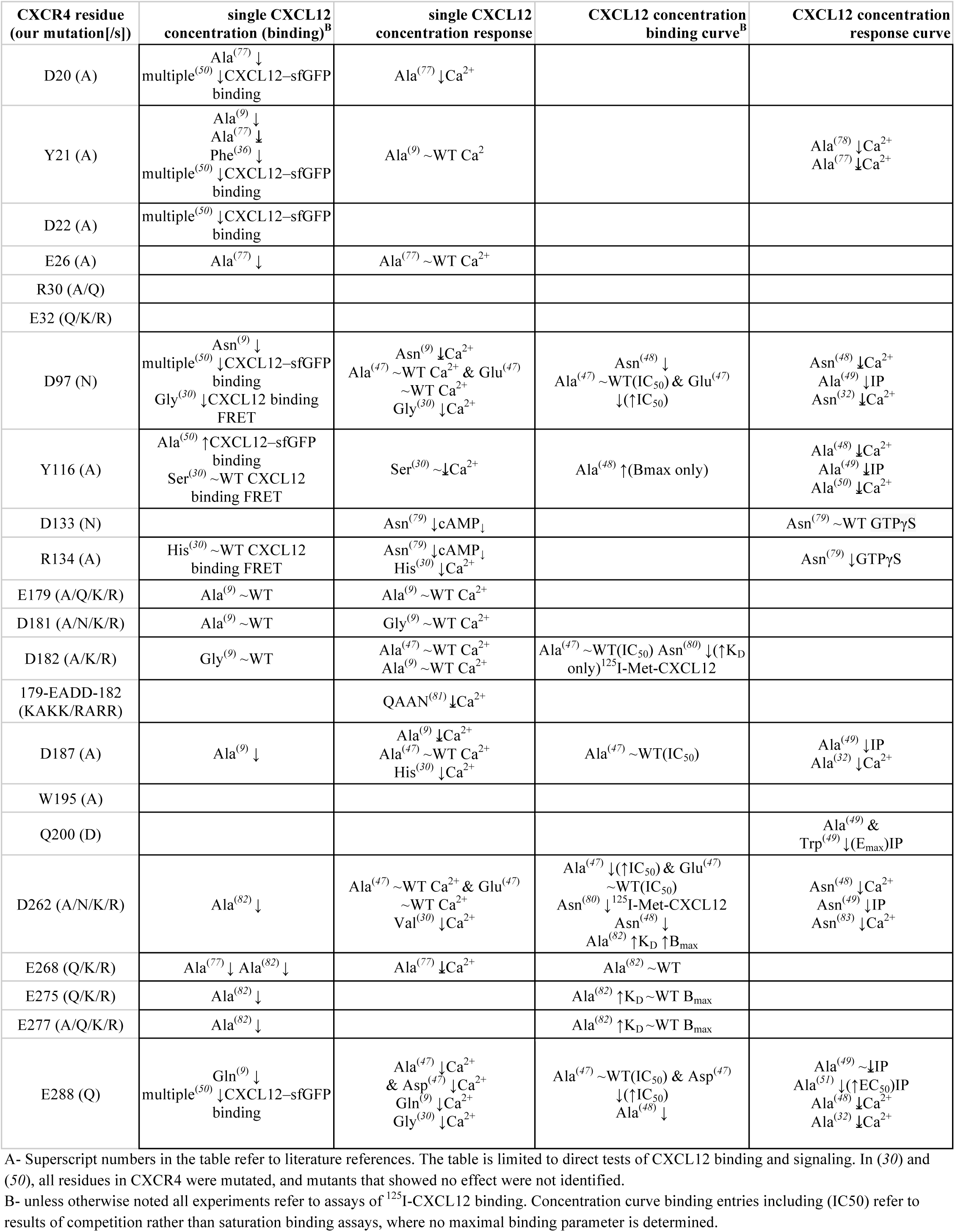
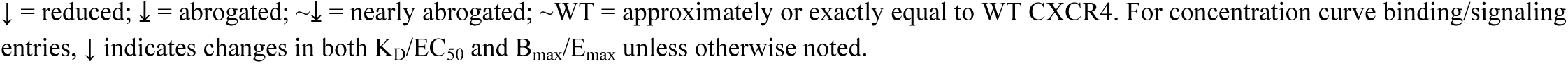
Existing literature mutagenesis effects on the CXCR4 residues tested in this study^A^.

Here we undertook a quantitative and systematic approach. Initially, CXCL12-induced β-arrestin-2 recruitment to the receptor mutants was monitored by a bioluminescence resonance energy transfer (BRET)-based assay (fig. S1A, *39, 46*). Select mutants with substantial effects on β-arrestin-2 recruitment were then characterized in a BRET-based mini-G_αi_ association assay (fig. S1B). For each mutant tested in either β-arrestin-2 or G_αi_ experiments, a full chemokine concentration response curve was generated, and both EC_50_ (potency) and E_max_ (efficacy) signaling parameters were obtained (Table 3). These direct BRET-based interaction assays are not subject to second messenger signal amplification, in contrast with the commonly employed calcium (Ca^2+^) mobilization and inositol monophosphate (IP) accumulation experiments (fig. S1C).

**Table 3.** Signaling parameters obtained in β arrestin-2 and mini-G_αi_ association BRET experiments.

| Mutation/truncation | *$\beta$ arrestin-2 association* | | *mini-$G_{ai}$ association* | | Bias factor ($\beta$) |
| --- | --- | --- | --- | --- | --- |
| | $\Delta pEC_{50} \pm SE$ | $E_{max} \pm SE$ | $\Delta pEC_{50} \pm SE$ | $E_{max} \pm SE$ | |
| $\Delta 1-7$ | $-0.06 \pm 0.08$ | $74 \pm 3^*$ | $-0.64 \pm 0.15^*$ | $94 \pm 6$ | $-0.46 \pm 0.17$ |
| $\Delta 1-10$ | $-0.02 \pm 0.06$ | $77 \pm 2^*$ | $-0.73 \pm 0.22^*$ | $83 \pm 9$ | $-0.67 \pm 0.24$ |
| $\Delta 1-15$ | $0.26 \pm 0.15$ | $30 \pm 2^*$ | $-1.7 \pm 0.15^*$ | $44 \pm 5^*$ | $-1.78 \pm 0.22$ |
| $\Delta 1-19$ | $0.05 \pm W$ | $14 \pm 4^*$ | | | |
| $\Delta 1-25$ | $-0.03 \pm W$ | $21 \pm 2^*$ | $-1.78 \pm 0.17^*$ | $27 \pm 4$ | |
| D20A | $-0.05 \pm 0.13$ | $83 \pm 5^*$ | | | |
| Y21A | $-0.59 \pm 0.08^*$ | $103 \pm 5$ | | | |
| D22A | $-0.53 \pm 0.07^*$ | $100 \pm 3$ | | | |
| E26A | $-0.13 \pm 0.07^*$ | $82 \pm 3^*$ | | | |
| 20-AAASMAA-26 | $-0.68 \pm 0.07^*$ | $90 \pm 3^*$ | $-0.83 \pm 0.29^*$ | $115 \pm 18$ | $-0.03 \pm 0.31$ |
| R30A | $-0.13 \pm 0.08^*$ | $49 \pm 2^*$ | | | |
| R30Q | $-0.07 \pm 0.09$ | $59 \pm 3^*$ | $-0.38 \pm 0.12^*$ | $70 \pm 3^*$ | $-0.23 \pm 0.16$ |
| E32Q | $0.05 \pm 0.08$ | $92 \pm 3^*$ | | | |
| E32K | $-0.02 \pm 0.05$ | $85 \pm 2^*$ | | | |
| E32R | $-0.12 \pm 0.06^*$ | $81 \pm 2^*$ | | | |
| D262N | $-0.59 \pm 0.06^*$ | $55 \pm 2^*$ | | | |
| D262A | $-0.05 \pm W$ | $20 \pm 2^*$ | | | |
| D262K | $-1.17 \pm 0.11^*$ | $22 \pm 2^*$ | $-1.59 \pm 0.11^*$ | $32 \pm 2^*$ | $-0.24 \pm 0.17$ |
| D262R | $-1.08 \pm 0.19^*$ | $24 \pm 3^*$ | | | |
| E268Q | $0.1 \pm 0.06$ | $120 \pm 4^*$ | | | |
| E268K | $0.07 \pm 0.05$ | $116 \pm 3^*$ | | | |
| E268R | $0.14 \pm 0.08$ | $103 \pm 4$ | | | |
| E275Q | $0.1 \pm 0.13$ | $93 \pm 5$ | | | |
| E275K | $-0.01 \pm 0.06$ | $104 \pm 3$ | | | |
| E275R | $0.26 \pm 0.08^*$ | $93 \pm 3^*$ | | | |
| E277A | $-0.21 \pm 0.09$ | $112 \pm 5^*$ | | | |
| E277Q | $0.05 \pm 0.06$ | $97 \pm 3$ | | | |
| E277K | $-0.47 \pm 0.05^*$ | $74 \pm 2^*$ | | | |
| E277R | $-0.51 \pm 0.08^*$ | $72 \pm 3^*$ | $-0.68 \pm 0.17^*$ | $72 \pm 6^*$ | $-0.17 \pm 0.2$ |
| E179A | $0.04 \pm 0.08$ | $104 \pm 4$ | | | |
| E179Q | $0.11 \pm 0.07$ | $94 \pm 3^*$ | | | |
| E179K | $0 \pm 0.09$ | $92 \pm 4^*$ | | | |
| E179R | $-0.05 \pm 0.07^*$ | $98 \pm 3$ | | | |
| D181A | $-0.17 \pm 0.09$ | $88 \pm 4^*$ | | | |
| D181N | $0 \pm 0.07$ | $89 \pm 3^*$ | | | |
| D181K | $-0.21 \pm 0.16$ | $77 \pm 6^*$ | | | |
| D181R | $-0.14 \pm 0.08$ | $69 \pm 3^*$ | | | |
| D182A | $0.02 \pm 0.11$ | $112 \pm 5^*$ | | | |
| D182K | $0.01 \pm 0.1$ | $85 \pm 4^*$ | | | |
| D182R | $0.01 \pm 0.05$ | $80 \pm 2^*$ | | | |
| 179-KAKK-182 | $-0.16 \pm 0.11^*$ | $39 \pm 2^*$ | $-1.03 \pm 0.15^*$ | $40 \pm 3^*$ | $-0.86 \pm 0.19$ |
| 179-RARR-182 | $-0.06 \pm 0.09$ | $25 \pm 1^*$ | | | |
| D97N | NR | NR |  |  |  |
| Y116A | NR | NR |  |  |  |
| D187A | $-0.37 \pm 0.23$ | $13 \pm 1^*$ | $-0.97 \pm 0.22^*$ | $33 \pm 4^*$ | $-0.2 \pm 0.32$ |
| E288Q | NR | NR |  |  |  |
| W195A | $-0.21 \pm 0.16^*$ | $48 \pm 3^*$ | $-0.52 \pm 0.23^*$ | $43 \pm 4^*$ | $-0.36 \pm 0.29$ |
| Q200D | $-0.08 \pm 0.08$ | $71 \pm 3^*$ | $-0.17 \pm 0.19$ | $61 \pm 4^*$ | $-0.15 \pm 0.21$ |
| D133N | $0.34 \pm 0.09^*$ | $52 \pm 2^*$ | $-0.24 \pm 0.16$ | $14 \pm 1^*$ | $-1.13 \pm 0.19$ |
| R134A | $0.32 \pm 0.16^*$ | $113 \pm 7^*$ | $-0.52 \pm 0.37$ | $6 \pm 1^*$ | $-2.13 \pm 0.41$ |

\* = significantly different from WT CXCR4 according to the Akaike's informative criteria (AICc) test; W - wide (poor response prevented precise EC<sub>50</sub> parameter calculation); NR = No analyzable response

To enable quantitative comparisons between WT and mutants, we designed the BRET experiments so that the observed potency, efficacy, and fraction of receptor on the cell surface did not vary within a substantial range of total WT CXCR4 expression levels (fig. S2A-E). Total expression was monitored for all mutants throughout the signaling experiments, and transfections were adjusted as needed to keep mutant expression within this range, though we note that only one perturbation required adjustment (fig. S2F, fig. S3A). To identify mutation-induced changes in receptor expression specifically at the cell surface, we independently determined both the total and surface expression levels for all but two CXCR4 mutants that demonstrated substantial effects in BRET experiments (any significant efficacy impairment of >15% and/or any significant potency impairment >0.1 log_10_ units) (fig. S3A-F). Throughout the results, we note all cases in which mutations impaired the fractional surface expression of CXCR4 (fig. S3C&F). For these mutants, no simple method was available to ensure surface expression comparable to WT, because the fraction of CXCR4 expressed at the cell surface did not vary in correspondence with total expression (fig. S2E).

In the sections that follow we systematically characterize selected mutants across the various CXCR4-CXCL12 interaction epitopes in a model-guided manner. Even though most of the residues mutated here have been probed in earlier studies (Table 2), a mutational study of CXCR4 has never been done in such a uniform quantitative way.

### Charge swap mutagenesis validates the predicted geometry of the CXCR4-CXCL12 complex

To validate the overall architecture and key polar interactions in the model, we used a strategy of “charge swap” mutagenesis. This strategy is superior to traditional single-sided loss-of-function mutagenesis because it can generate information about direct pairwise residue contacts between the receptor and the chemokine via functional rescue.

Our CXCR4-CXCL12 model suggests that Asp^262^(6.58) of CXCR4 forms a CRS2-anchoring ionic interaction with Arg^8^ of CXCL12 (Fig. 2A). Consistent with the model, D262A, D262K and D262R mutations all nearly abolished CXCL12-mediated β-arrestin-2 recruitment, while D262N showed a lesser but still severe effect (Fig. 2B). D262K also severely impaired G_αi_ recruitment to CXCR4 (Fig. 2C). Another important intermolecular salt bridge predicted by the model is made between CXCR4 Glu^277^(7.28) and Arg^12^ of CXCL12 (Fig. 2A). Mutation of Glu^277^(7.28) to Ala or Gln had no negative effect on signaling, whereas E277K and E277R both showed potency deficits and an approximately 30% decrease in β-arrestin-2 recruitment efficacy (Fig. 2D, Table 3). CXCR4(E277R) demonstrated similar potency and efficacy impairment in the G_αi_ assay (Fig. 2C). By contrast, mutations of neighboring acidic residues (fig. S4A) Glu^268^(ECL3) and Glu^275^(7.26) to Gln, Lys, and Arg produced almost no negative effect (fig. S4B&C); in fact we observed slight but significant increases in efficacy for CXCR4(E268Q/K) and potency in the case of CXCR4(E275R) (fig. S4C, Table 3).

**Fig. 2.**
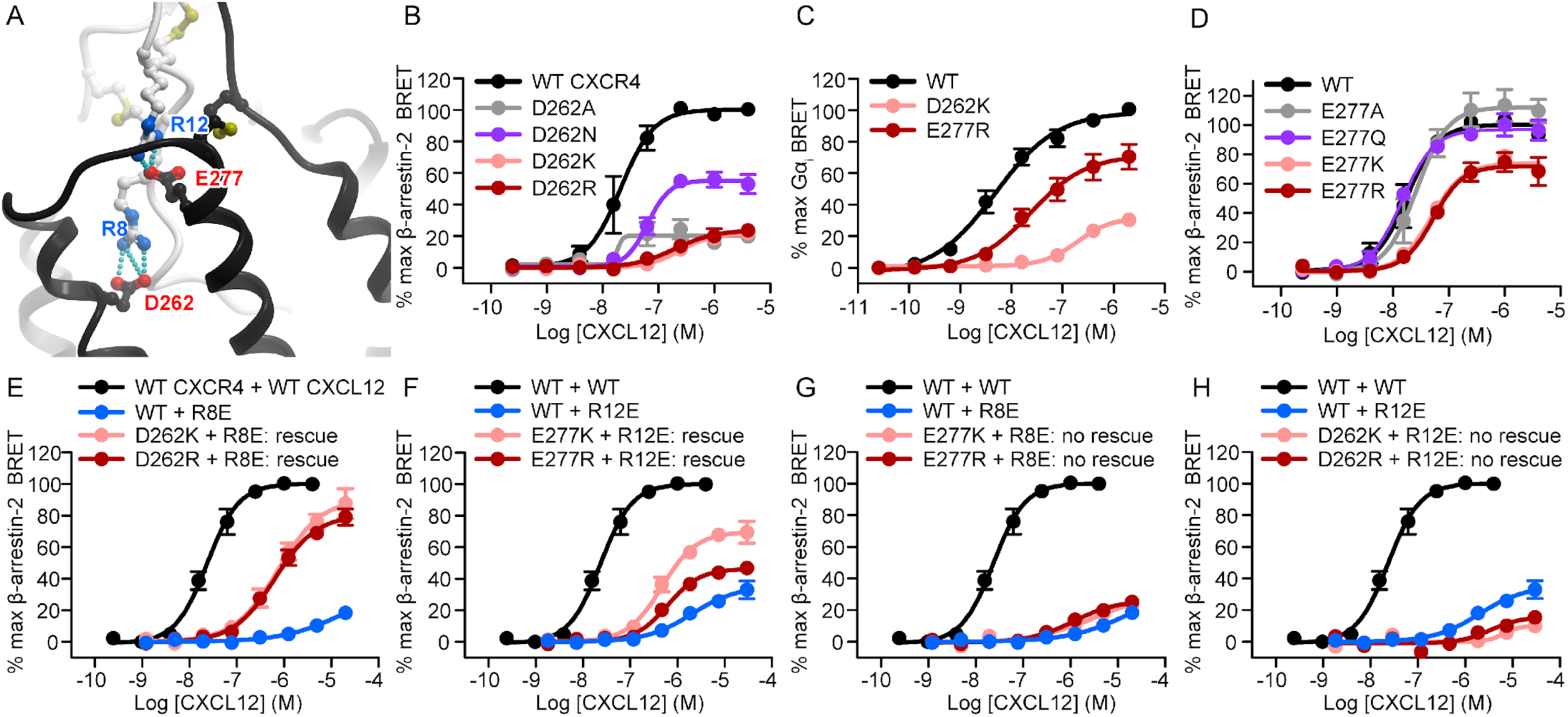
Reciprocal charge reversal experiments validate the model geometry and establish the roles and the interaction partners of CXCL12 residues Arg^8^ and Arg^12^. (**A**) The two salt bridges proposed by the model to determine the orientation of CXCL12 with respect to CXCR4 involve CXCL12 Arg^8^ and Arg^12^ paired with CXCR4 Asp^262^(6.58) and Glu^277^(7.28), respectively The chemokine and the receptor are shown in white and black ribbon, respectively. (**B**) β-arrestin-2 association BRET ratio data for a series of CXCR4 Asp^262^(6.58) mutants (D262A/N/K/R) after stimulation with varying CXCL12 concentrations for 20 minutes. (**C**) Mini-G_ɑi_ association BRET ratio data for CXCR4(D262K) and CXCR4(E277R) after stimulation with varying CXCL12 concentrations for 1 minute. (**D**) β-arrestin-2 CXCL12 concentration-response data obtained for a series of CXCR4 Glu^277^(7.28) mutants (E277A/Q/K/R). (**E**) β-arrestin-2 CXCL12 concentration-response data obtained by stimulating WT CXCR4 with WT CXCL12, WT CXCR4 with CXCL12(R8E), CXCR4(D262K) with CXCL12(R8E), and CXCR4(D262R) with R8E CXCL12. (**F**) CXCL12 concentration-response β-arrestin-2 BRET data for WT CXCR4 stimulated with WT CXCL12, WT CXCR4 with CXCL12(R12E), CXCR4(E277K) with CXCL12(R12E), and CXCR4(E277R) with CXCL12(R12E). (**G**) β-arrestin-2 CXCL12 concentration-response data obtained by stimulating WT CXCR4 with WT CXCL12, or by stimulating either WT CXCR4, CXCR4(E277K), or CXCR4(E277R) with CXCL12(R8E). (**H**) β-arrestin-2 CXCL12 concentration-response data obtained by stimulating WT CXCR4 with WT CXCL12, or by stimulating either WT CXCR4, CXCR4(ED262K), or CXCR4(ED262R) with CXCL12(R12E). In **B**-**H**, data represent the mean values from at least three independent experiments, each performed in duplicate with data normalized to the E_max_ of WT CXCR4 (WT CXCR4 + WT CXCL12 for charge swap experiments) tested in the same experiments. The same WT CXCR4 + WT CXCL12, WT CXCR4 + CXCL12(R8E), and WT CXCR4 + CXCL12(R12E) datasets are shown in panels **E**/**G** and **F**/**H**. Error bars indicate SEM. For this and all subsequent figures, error bars that are smaller than the circle visualizing the mean are not shown.

On the chemokine side, an R8E mutation of CXCL12 practically eliminated its ability to activate CXCR4 (Fig. 2E). However, applying CXCL12(R8E) to cells expressing CXCR4(D262K/R) led to robust activation, with efficacy that by far exceeded that of the same mutants tested individually (Fig. 2E). In fact the efficacy approached that observed for WT CXCR4-CXCL12 signaling. A CXCL12 R12E mutation also greatly decreased the potency and efficacy of CXCR4 activation (Fig. 2F), although not to the same extent as R8E (Fig. 2E). But again, CXCR4(E277K) rescued the signaling of CXCL12(R12E) substantially, as did CXCR4(E277R) (although with a lower efficacy) (Fig. 2F). The functional rescue effects were specific, as no rescue was observed when chemokine mutants from each of the two predicted salt bridges were combined with receptor mutants from the other salt bridge (namely CXCL12(R12E) with CXCR4(D262K/R) or CXCL12(R8E) with CXCR4(E277K/R), Fig. 2G&H).

We note that the potency of either rescuing combination did not reach that of the WT receptor-WT chemokine combination. Furthermore, in the case of CXCR4(E277K/R)-CXCL12(R12E) the rescue was not reciprocal, as the efficacy exceeded that of WT CXCR4-CXCL12(R12E) but not that of CXCR4(E277K/R)-WT CXCL12. Nevertheless, the fact that receptor mutations restored the signaling deficits of the chemokine mutations indicates that the corresponding residues are in direct contact in the complex. The inability to completely restore signaling to WT levels can be attributed to the complexity of the interface where other residue interactions play a role, or to the requirement of a precise spatial arrangement of residues for full signaling capacity. In fact, the salt bridge between CXCR4 Glu^277^(7.28) and CXCL12 Arg^12^ is part of a larger interconnected network of hydrogen bonding interactions also involving CXCR4 Arg^30^(NT) (Fig. 3A), a residue at the junction of the receptor N-terminus and TM1 that is predicted to be involved in coordination of CRS1.5 interactions. In our experiments, even a conservative substitution of Arg^30^(NT) with Gln resulted in a substantial efficacy loss of approximately 40% and 30% in β-arrestin-2 and G_αi_ experiments respectively, while the alanine mutant CXCR4(R30A) showed a significant potency impairment as well (Fig. 3B-C, Table 3).

**Fig. 3.**
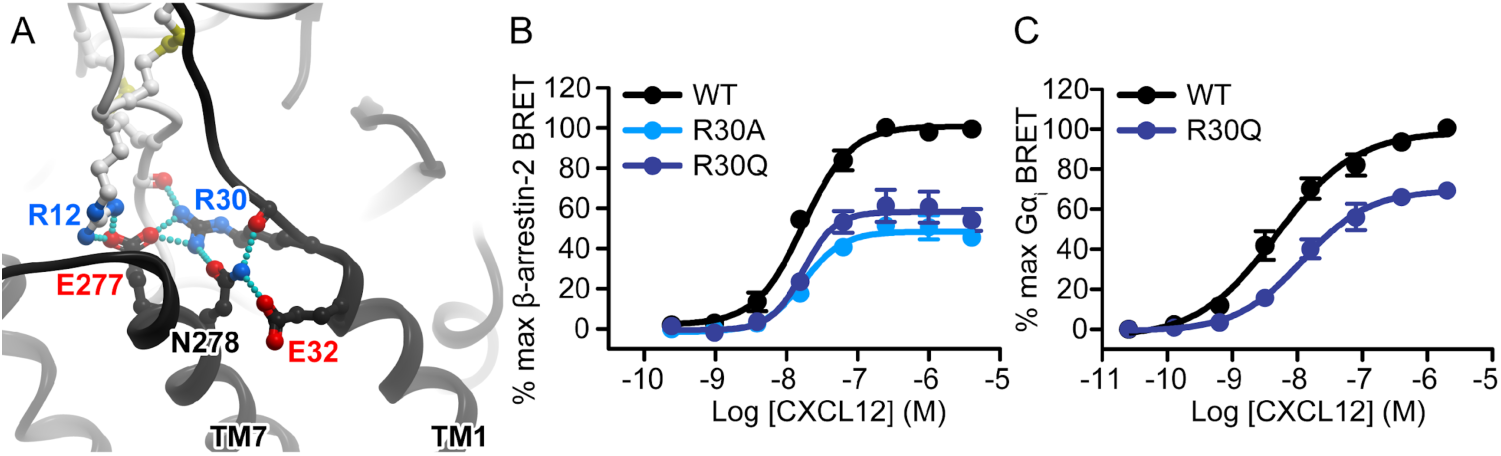
CXCR4 residue Arg^30^(NT) is an important mediator of CXCL12 signaling towards both G_ɑi_ and β-arrestin-2. (**A**) The model predicts CXCR4 Arg^30^(NT) to be the central residue in the CRS1.5 hydrogen-bonding network between CXCR4 and CXCL12. (**B**) β-arrestin-2 association BRET ratio data for CXCR4(R30A) and CXCR4(R30Q) mutants after stimulation with varying CXCL12 concentrations for 20 minutes. (**C**) Mini-G_ɑi_ association BRET ratio data for CXCR4(R30Q) after stimulation with varying CXCL12 concentrations for 1 minute. All data represent the mean values from at least three independent experiments, each performed in duplicate with data normalized to the E_max_ of WT CXCR4 tested in the same experiments. Error bars indicate SEM.

We also applied charge swap mutagenesis to probe an alternative geometry of the CXCR4-CXCL12 complex developed by Ziarek and colleagues (*28*). In that model, Arg^8^ of CXCL12 is purported to interact directly with CXCR4 Glu^32^(NT) rather than with Asp^262^(6.58), and CXCL12 Arg^12^ is predicted to interact with CXCR4 Asp^181^(ECL2) rather than with Glu^277^(7.28). When tested in the β-arrestin-2 assay, Gln, Lys, and Arg mutations of CXCR4 Glu^32^(NT) produced modest efficacy impairments (8, 15, 19% reductions, respectively), and CXCR4(E32R) produced a significant potency impairment as well (Fig. 4A, Table 3). However, we observed no rescue of the severely impaired signaling of CXCL12(R8E) when combined with CXCR4(E32K/R) (Fig. 4B). Similarly we observed no rescue of function when either CXCR4(D181K) or CXCR4(D181R) were combined with CXCL12(R12E) (Fig. 4C). These findings argue against the geometry of the complex proposed by Ziarek and colleagues (*28*), but are consistent with our model, where CXCR4 Glu^32^(NT) is 15.6 Å away from CXCL12 Arg^8^ (Cα atom distance), and CXCR4 Asp^181^(ECL2) is on the opposite side of the CXCR4 binding pocket from Glu^277^(7.28) and 16.5 Å from CXCL12 Arg^12^ (Fig. 4D). The discrepancy is due to a different rotational position of the chemokine relative to the receptor in the two models.

**Fig. 4.**
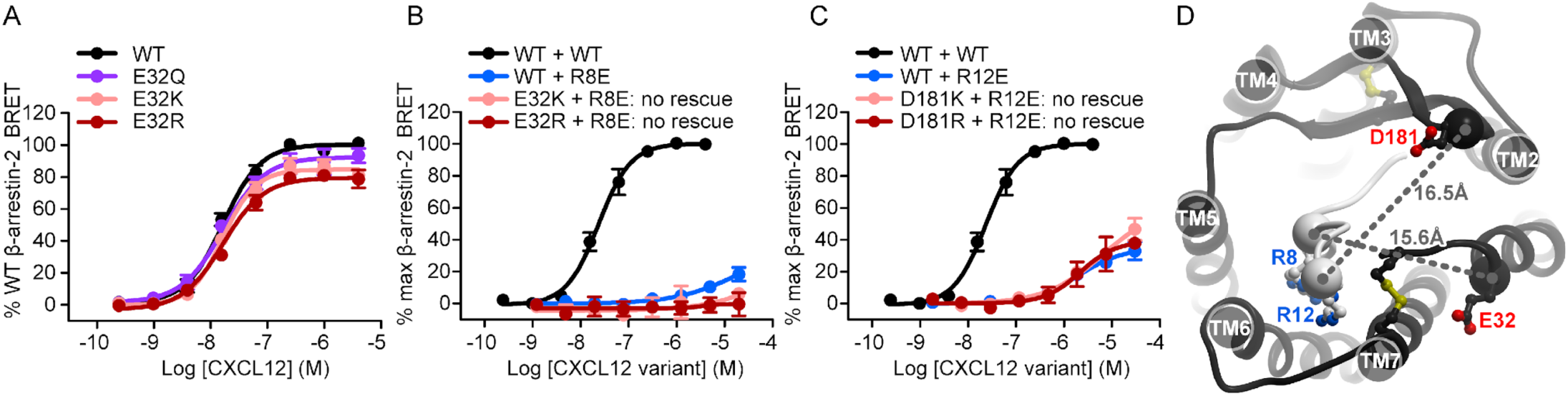
Charge swap experiments do not support the pairing of CXCR4 residues Glu^32^(NT) and Asp^181^(ECL2) with CXCL12 Arg^8^ and Arg^12^, respectively. (**A**) β-arrestin-2 association BRET ratio data for a series of CXCR4 Glu^32^(NT) mutants (E32Q/K/R) after stimulation with varying CXCL12 concentrations for 20 minutes. (**B**) β-arrestin-2 CXCL12 concentration-response data obtained by stimulating WT CXCR4 with WT CXCL12, or by stimulating either WT CXCR4, CXCR4(E32K), or CXCR4(E32R) with CXCL12(R8E). (**C**) β-arrestin-2 CXCL12 concentration-response data obtained by stimulating WT CXCR4 with WT CXCL12, or by stimulating either WT CXCR4, CXCR4(D181K), or CXCR4(D181R) with CXCL12(R12E). All data represent the mean values from at least three independent experiments, each performed in duplicate with data normalized to the E_max_ of WT CXCR4 tested in the same experiments. Error bars indicate SEM. The same WT CXCR4 + WT CXCL12, WT CXCR4 + CXCL12(R8E), and WT CXCR4 + CXCL12(R12E) data are shown in (**B**) and (**C**) as those shown in (**E**-**H**) of Figure 1. (**D**) The relative location of the two control residue pairs, CXCR4 Asp^181^(ECL2) and CXCL12 Arg^12^, and CXCR4 Glu^32^(NT) and CXCL12 Arg^8^, in the model of the CXCR4-CXCL12 complex.

### The N-terminus of CXCR4, including CRS0.5, contributes to CXCR4 signaling efficacy

Although all three receptor-chemokine complexes crystallized thus far utilize full length receptors, the solved structures lack density for a large stretch of residues in the receptor N-termini (residues 1-22 in CXCR4). For the vMIP-II-bound structure of CXCR4 (*14*), the visible density contains only the proximal N-terminus (residues 23-27) interacting with the N-loop/40s loop groove of the chemokine, and provides no information for the role of any of the putative sulfotyrosines (sTyr^7^, sTyr^12^, sTyr^21^). As described above, our model suggests that the entire receptor N-terminus engages CXCL12 by wrapping around the chemokine globular domain, with the distal N-terminus (CRS0.5 residues 3-GISIsY-7) forming an anti-parallel β-sheet with the β_1_-strand of the chemokine (25-HLKIL-29). In order to globally probe the functional role of the CRS0.5/CRS1 interaction, we generated several CXCR4 constructs with truncations of 7, 10, 15, 19, and 25 residues (Fig. 5A). Successively longer deletions produced progressively larger reductions in β-arrestin-2 recruitment efficacy, with CXCR4(Δ1-19) and CXCR4(Δ1-25) displaying <25% WT efficacy (Fig. 5B, Table 3). Moreover, the efficacy impairment of CXCR4(Δ1-15) (30% WT efficacy remaining) was almost the same as that of a 10-residue longer truncation, CXCR4(Δ1-25), indicating that it is the distal and not the proximal N-terminus that plays a dominant role in the signaling efficacy of the receptor. The truncations produced no observable impairments in β-arrestin-2 recruitment *potency*, although CXCR4(Δ1-19) and CXCR4(Δ1-25) yielded such poor signaling that their EC_50_ values could not be accurately fitted. Note that despite the size and location of the truncations, only CXCR4(Δ1-19) showed notably reduced surface expression (65% WT), but not to any extent that would explain its severely impaired (<15% WT) signaling efficacy (fig. S3B-C).

**Fig. 5.**
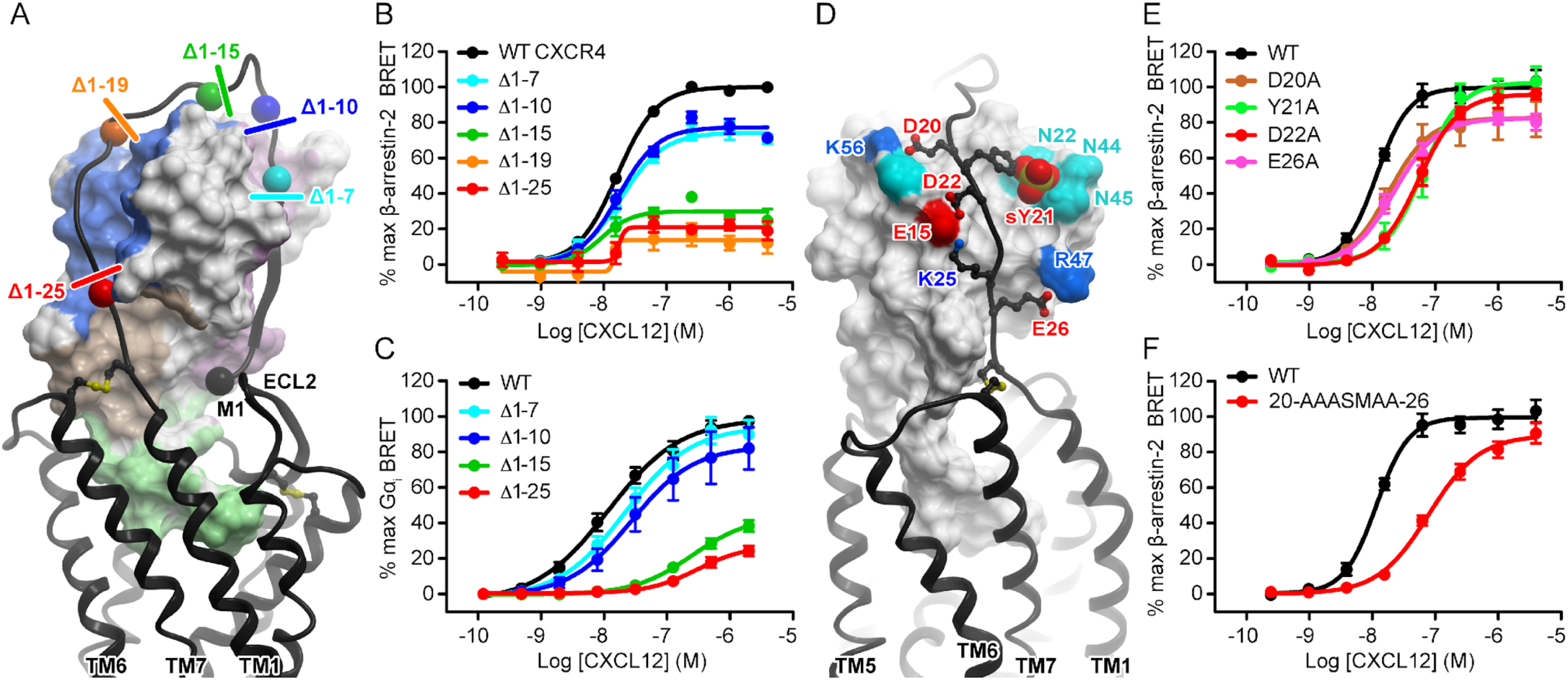
The N-terminus of CXCR4 is important for both the efficacy and potency of CXCR4-CXCL12 signaling. (**A**) Progressive N-terminal truncations of CXCR4 illustrated in the context of the model of the CXCR4-CXCL12 complex. β-arrestin-2 association BRET ratio data for a series of CXCR4 N-terminal truncations (Δ1-7, Δ1-10, Δ1-15, Δ1-19, Δ1-25) after stimulation with varying CXCL12 concentrations for 20 minutes. (**C**) Mini-G_ɑi_ association BRET ratio data for a series of CXCR4 N-terminal truncations (Δ1-7, Δ1-10, Δ1-15, Δ1-25) after stimulation with varying CXCL12 concentrations for 1 minute. (**D**) The predicted polar interactions between the proximal N-terminus of CXCR4 and the N-loop/40s loop groove of CXCL12. (**E**) β-arrestin-2 CXCL12 concentration-response data for CXCR4(D20A), CXCR4(Y21A), CXCR4(D22A), and CXCR4(E26A), or (**F**) for CXCR4(20-AAASMAA-26), in which in which all four CXCR4 residues tested in (**E**) along with Lys^25^(NT) are mutated to Ala simultaneously. All data represent the mean values from at least three independent experiments, each performed in duplicate with data normalized to the E_max_ of WT CXCR4 tested in the same experiments. Error bars indicate SEM.

When examined in the G_αi_ BRET assay, the same overall pattern of reduced efficacy with increasing truncation length was observed (Fig. 5C), although in this case the 7- and 10-residue truncations are not significantly impaired in efficacy compared to WT CXCR4 (Table 3). In contrast to the β-arrestin recruitment assay, we observed major reductions in potency with progressive receptor truncations in the G_αi_ association experiments (Fig. 5C). In order to ensure the results of N-terminal truncations were not in some way related to the presence of the N-terminally fused HA-tag, we removed the tag in the case of mini-G_αsi_ BRET experiments, and while the signaling results were quite similar (Fig. 5C) to those of β-arrestin-2 experiments, we noted a greater negative impact of the truncations on receptor surface expression (fig. S3E-F) without the HA tag present. Because the CXCR4(Δ1-15) and CXCR4(Δ1-25) truncations showed 40% and 33% WT surface expression respectively in the G_αi_ assays (fig. S3F), the apparent efficacy impairments (44 and 27% WT efficacy remaining) must be interpreted with caution.

Charge complementarity, suggested by the model, likely defines the position of the receptor N-terminus as it contacts residues from the N-loop, the 40s loop, and the C-terminal helix of the chemokine, culminating with CRS0.5. Specifically, CXCR4 Glu^26^(NT), Lys^25^(NT), Asp^22^(NT) and Asp^20^(NT) interact with CXCL12 Arg^47^ (40s loop), Glu^15^ and His^17^ (N-loop), and Lys^56^ while the sulfated CXCR4 Tyr^21^(NT) binds CXCL12 Asn^22^ (3_10_ helix), Asn^44^ and Asn^45^ (40s loop) (Fig. 5D). Accordingly, in addition to N-terminal truncations of CXCR4, we also examined alanine mutants of the above charged residues. CXCR4(D20A) and CXCR4(E26A) mutations produced small (<20%) but significant efficacy impairments while CXCR4(sY21A), CXCR4(D22A), and CXCR4(E26A) mutations all produced significant potency impairments (Fig. 5E, Table 3). When all of the alanine mutations were combined, the potency defect was not much greater than that of either CXCR4(sY21A) or CXCR4(D22A) alone and only a small (10%) efficacy decrease was observed (Fig. 5F, fig. S5, Table 3), consistent with the traditional view of CRS1 as a region that principally contributes binding affinity to the CXCR4-CXCL12 complex. However, it appears that multiple N-terminal residues function in unison rather than any individual residue dominating the contribution. Taken together with the results of the truncation mutants, these data challenge the established view of the receptor N-terminus as only an affinity determinant and reveal that it also plays a role in signaling efficacy, but without any apparent “hotspot residues”.

### Negatively charged residues in the CXCR4 ECL2 hairpin are important for both β-arrestin-2 and G_ɑi_ recruitment efficacy

The extracellular loop 2 (ECL2) of CXCR4 forms a β-hairpin whose tip contains three negatively charged residues (Glu^179^(ECL2), Asp^181^(ECL2), and Asp^182^(ECL2)) that in our model, are in proximity to a positively charged patch of the chemokine involving CXCL12 residues His^25^, Lys^27^ and Arg^41^ (Fig. 6A&B). To probe the role of this charge cluster, we mutated the constituent residues, both separately and simultaneously. Most single point mutations of the ECL2 hairpin (E179A/Q/K/R, D181A/N/K, and D182A/K/R) produced modest (25% or less) but significant efficacy reductions with the exception of D181R which showed a 30% reduction in maximal β-arrestin-2 recruitment, and only one mutation (E179R) had a modest but significant effect on potency (Fig. 6C-E, Table 3). However, when Glu^179^(ECL2), Asp^181^(ECL2), and Asp^182^(ECL2) were all mutated to either lysine (179-KAKK-182) or arginine (179-RARR-182), β-arrestin-2 recruitment was reduced to just 39% and 25% of WT efficacy, respectively, with 179-KAKK-182 displaying a substantial potency impairment as well (Fig. 6F, Table 3). In the G_αi_ BRET assay, 179-KAKK-182 also displayed a large reduction in efficacy, along with a much larger decrease in potency than was observed in β-arrestin-2 recruitment (Fig. 6G, Table 3). This likely indicates the lack of specific intermolecular contacts involving these residues, whereas the overall negative charge of the ECL2 hairpin is important for favorable electrostatic attraction to the major basic surface of the chemokine.

**Fig. 6.**
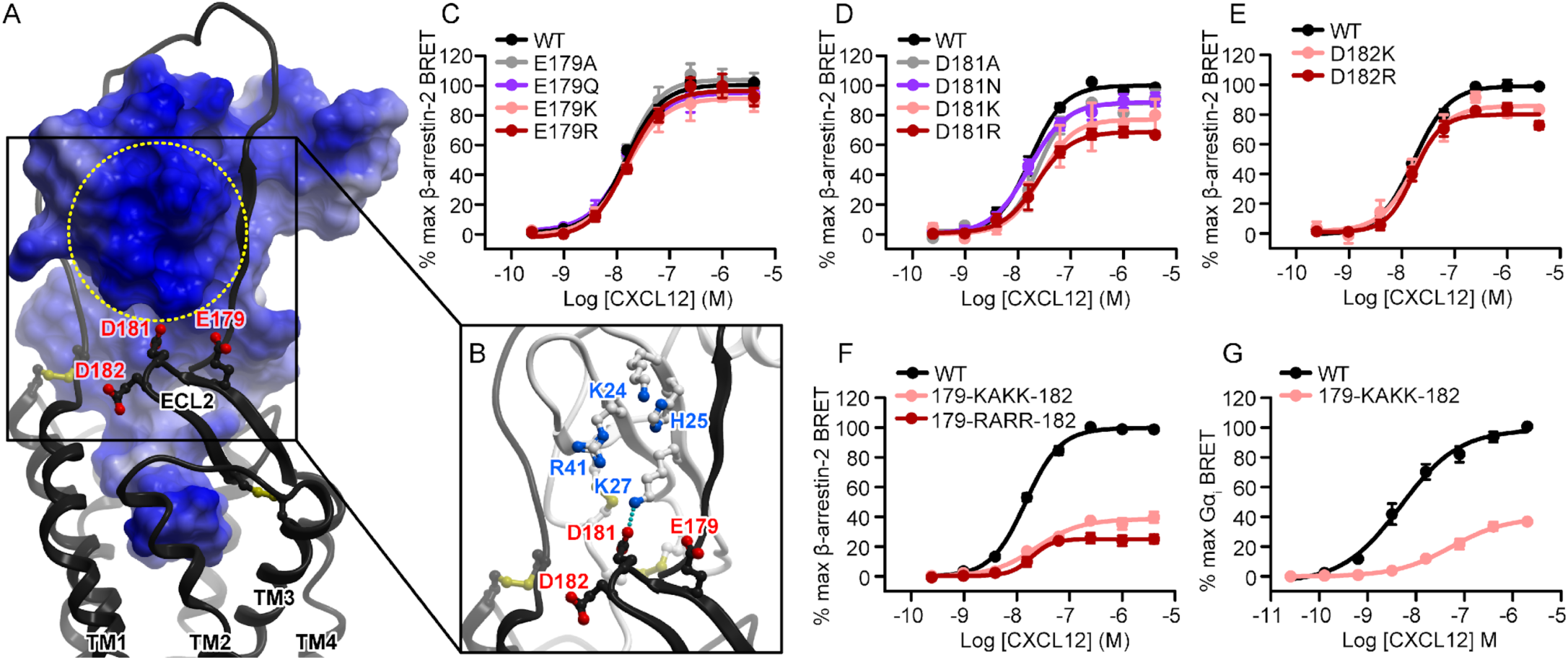
The ECL2 hairpin loop is synergistically important for CXCR4 activation. (**A**-**B**) The negatively charged β-hairpin of CXCR4 ECL2 (residues 179-EADD-182) is predicted to be proximal to the basic patch on the CXCL12’s three-stranded β-sheet. In (**A**), the receptor is shown as a ribbon and the chemokine as a surface mesh colored blue-to-red by electrostatic potential. The entire chemokine surface is basic, while the indicated patch stands out because of a higher-than-average concentration of positively charged residues. In (**B**), the chemokine is shown as a ribbon and basic residues forming the patch are indicated. (**C**-**F**) β-arrestin-2 association BRET ratio data for (**C**) a series of Glu^179^(ECL2) mutations (E179A/Q/K/R), (**D**) a series of Asp^181^(ECL2) mutations (D181A/N/K/R), (**E**) a pair of Asp^182^(ECL2) mutations (E182K/R), and (**F**) CXCR4(179-KAKK-182) and CXCR4(179-RARR-182), in which all CXCR4 residues tested in (**C**-**E**) are mutated to Lys or Arg simultaneously, after stimulation with varying CXCL12 concentrations for 20 minutes. (**G**) Mini-Gɑ_i_ association BRET ratio data for CXCR4(179-KAKK-182) after stimulation with varying CXCL12 concentrations for 1 minute. All data represent the mean values (with error bars indicating SEM) from at least three independent experiments, each performed in duplicate with data normalized to the E_max_ of WT CXCR4 tested in the same experiments.

### CXCR4 residues directly contacting the extreme CXCL12 N-terminus are critical for activation

CRS2 is the best characterized interaction epitope of the CXCR4-CXCL12 complex (*9, 30, 32, 47-51*), with numerous studies reporting deleterious effects of mutations of charged residues in the receptor binding pocket. This is likely because interactions between these residues and the flexible chemokine N-terminus are crucial for CXCL12-mediated CXCR4 activation. Here we sought to revisit these findings in the context of our 3D model, employing the amplification-free assays, quantitative approaches, and with rigorous surface expression monitoring.

In our model, CXCL12 Lys^1^ interacts with CXCR4 Asp^97^(2.63) through the N-terminal amine group, and with CXCR4 Glu^288^(7.39) through the Lys^1^ side chain (Fig. 7A, *14*). Individual mutations of D97N and E288Q eliminated CXCL12-mediated β-arrestin-2 recruitment despite being conservative substitutions (Fig. 7B). CXCR4 residue Tyr^116^(3.32) sits just underneath CXCL12 Pro^2^ and is thought to couple CXCR4-CXCL12 engagement to intracellular conformational changes within the receptor. In our hands, a Y116A mutation of CXCR4 abrogated β-arrestin-2 recruitment completely (Fig. 7B). Finally, in the model, CXCR4 residue Asp^187^(ECL2), near the base of the CXCR4 ECL2 hairpin, is positioned to interact with CXCL12 Tyr^7^ and the backbone amide of Val^3^. Consistent with this, CXCR4(D187A) was almost completely inactive (<15% WT efficacy) in β-arrestin-2 recruitment (Fig. 7B, Table 3), while some activation was preserved in the G_αi_ association experiments (approximately 30% WT) (Fig. 7C, Table 3). These data are consistent with the exceptional sensitivity of CXCR4 activation to residue substitutions in the binding pocket, which also mirrors the sensitivity of receptor activation to N-terminal modifications of CXCL12 (*2*). Combined with the structural model, the data provides insight into the initial steps of CXCR4 activation by CXCL12.

**Fig. 7.**
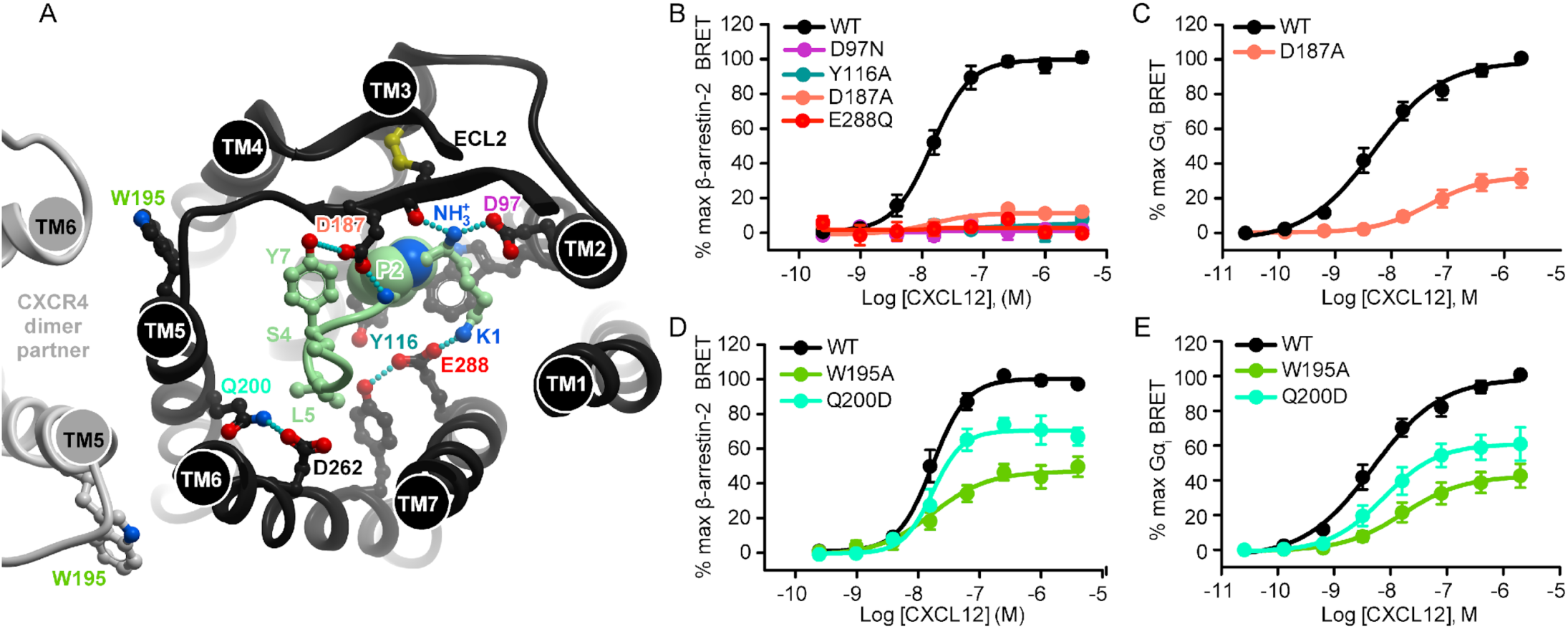
CRS2 receptor binding pocket mutations abrogate β-arrestin-2 recruitment. (**A**) The N-terminus of CXCL12 (light green) in the binding pocket of CXCR4. The receptor is viewed “top-down”, i.e. across the plane of the membrane from the extracellular side. Residues mentioned in the text are indicated. The hypothetical position of the CXCR4 dimer (based on the crystallographic dimer identified in PDB 3ODU) is shown on the left in light gray. (**B**) β-arrestin-2 association BRET ratio data for CXCR4(D97N), CXCR4(Y116A), CXCR4(D187A), and CXCR4(E288Q) after stimulation with varying CXCL12 concentrations for 20 minutes. (**C**) Mini-G_ɑi_ association BRET ratio data for CXCR4(D187A) after stimulation with varying CXCL12 concentrations for 1 minute. (**D**) β-arrestin-2 BRET or (**E**) Mini-G_ɑi_ BRET CXCL12 concentration-response data for CXCR4(W195A) and CXCR4(Q200D). All data represent the mean values (with error bars indicating SEM) from at least three independent experiments, each performed in duplicate with data normalized to the E_max_ of WT CXCR4 tested in the same experiments.

### Transmembrane helix 5 mutations cause impaired β-arrestin-2 recruitment through unknown mechanisms

Little attention has been paid to the role of residues in TM5 and the major subpocket of CXCR4 in prior mutagenesis studies of CXCR4-CXCL12 signaling, given the predominant role of minor pocket residues and sensitivity of CXCL12 N-terminus to mutations. This is challenged by our newly refined model of the complex, which features the CXCL12 N-terminal residues Ser^4^ and Leu^5^ in the major subpocket of CXCR4. Additionally, in our studies of the atypical chemokine receptor 3 (ACKR3), a homologous receptor that also binds CXCL12, we observed that the major subpocket residues Trp^208^(5.34) and Glu^213^(5.39) were important for CXCL12-mediated arrestin recruitment (*39*). We therefore tested the effects of mutating the corresponding CXCR4 residues, Trp^195^(5.34) and Gln^200^(5.39), on effector recruitment to CXCR4. CXCR4 Q200D and W195A mutations both reduced the efficacy of β-arrestin-2 and G_αi_ recruitment, and W195A also impaired potency (Fig. 7D-E). In our model, Gln^200^(5.39) mediates the TM5 interaction with TM6 via hydrogen bonding to Asp^262^(6.58), which itself is a key chemokine-coordinating residue (Fig. 7A). Moreover, in many GPCRs, activation is associated with a counter-clockwise rotation of TM5, which helps to shape the G protein binding pocket at the intracellular side (*52–55*). The proximity of Gln^200^(5.39) to CXCL12 Ser^4^ prompts a hypothesis that these two residues may engage in direct interaction, facilitating the TM5 rotation and an active-like conformation of the CXCR4 TM bundle. By contrast, Trp^195^(5.34) points away from the receptor core and is unlikely to engage in direct interaction with the chemokine. However, this residue has been proposed to mediate receptor dimerization (*56*), which may be a possible explanation for the effects of its mutation.

### Quantifying the apparent pathway bias in mutation-induced signaling impairments

Mutations of the ECL2 hairpin and the N-terminal CXCR4 truncations, but not other mutations tested, show unexpected discrepancies, as they display stronger potency impairment in G_αi_ experiments than in β-arrestin-2 experiments. Fortunately, our approach enabled quantification of the relative effects of mutants tested in the two assays using the equiactive comparison method of bias calculation (*45*). The N-terminal truncations and the ECL2 triple mutant E179K/D181K/D182K were markedly biased, all toward β-arrestin-2 recruitment (Fig. 8A). This is intriguing because of the general interest in GPCR biased signaling as a strategy for obtaining improved therapeutics and the desire to explain bias from the structural perspective. While the mechanistic basis for the biased results of these mutations and truncations remains unclear, it is notable that all bias-associated residues were found to interact with the same general region of the chemokine globular core (Fig. 8B). This finding recalls a prior report of signaling bias by dimeric CXCL12 (*41, 57*), which effectively prevents receptor interactions with the same residues in CXCL12 β1 strand (fig. S6, *38*).

**Fig. 8.**
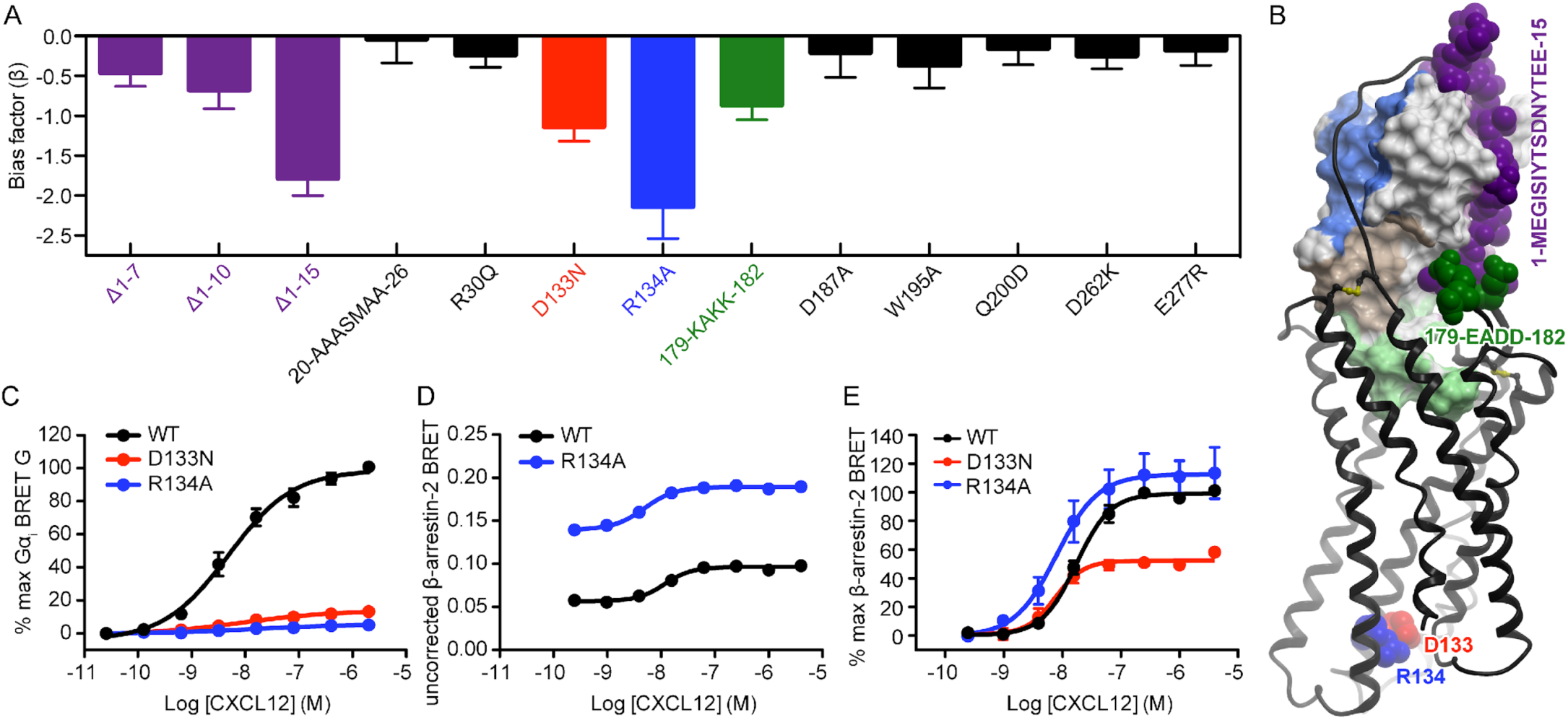
Mutations of CRS0.5 and ECL2 appear to cause biased signaling. (**A**) Equiactive bias factor (β) for all CXCR4 mutants that were tested in both β-arrestin-2 and mini-G_ɑi_ association BRET experiments. Error bars indicate combined errors of the EC_50_ and E_max_ values used to calculate β, and colored bars indicate that the absolute value of β > 2 times the combined error. A negative bias factor indicates bias towards β-arrestin-2. (**B**) When mapped onto the model of the CXCR4-CXCL12 complex, biased-associated residues at the chemokine interface cluster on the same side. The intracellular residues Asp^133^(3.49) and R^134^(3.50) are not in direct contact with the chemokine but are shown for reference. (**C**) Mini-G_ɑi_ association BRET ratio data for CXCR4(D133N) and CXCR4(R134A) after stimulation with varying CXCL12 concentrations for 1 minute. (**D**) Uncorrected β-arrestin-2 association BRET ratio data (without background signal subtraction or normalization) for CXCR4(R134A) after stimulation with varying CXCL12 concentrations for 20 minutes. Data from one independent experiment representative of three is shown. (**E**) β-arrestin-2 BRET CXCL12 concentration-response data for CXCR4(D133N) and CXCR4(R134A). In **C** and **E**, data represent the mean values (with error bars indicating SEM) from at least three independent experiments, each performed in duplicate with data normalized to the E_max_ of WT CXCR4 tested in the same experiments.

In order to calibrate the level of bias in the above mutants, we examined residues in the CXCR4-G protein interface. The conserved DRY motif in the intracellular region of TM3 participates directly in G protein coupling (*58*) and is known to modulate G protein signaling by class A GPCRs (*59, 60*). Mutation of the central Arg(3.50) residue of the DRY motif in other GPCRs (*61, 62*), including CCR5 (*63*), has been shown to produce receptor bias wherein G protein signaling is depleted but arrestin interactions are preserved or enhanced. Consistent with these findings, the CXCR4 R134A mutation almost completely abrogated G_αi_ recruitment (Fig. 8C). However, in β-arrestin-2 recruitment assays, CXCR4(R134A) showed not only increased constitutive association (Fig. 8D), but also higher efficacy and stronger potency in CXCL12-mediated activation (Fig. 8E). Although CXCR4(R134A) suffered from a slight surface expression deficit (fig. S3C,F), this decrease was not great enough to explain the observed major bias towards β-arrestin-2 vs G_αi_ recruitment; in fact, the decreased surface expression may reflect increased internalization resulting from the constitutive association with β-arrestin-2. The CXCR4(D133N) mutant showed an approximately 50% reduction in β-arrestin-2 recruitment efficacy along with a significant improvement in potency (Fig. 8E, Table 3), but similar to CXCR4(R134A), it displayed essentially no G_αi_ recruitment (Fig. 8C). CXCR4(D133N) was particularly deficient in surface expression in the β-arrestin-2 recruitment assay (fig. S3C), so its efficacy may be artificially impaired, though we note in such a case the true bias for this mutant toward β-arrestin-2 would in fact be higher than is apparent from the available data.

CXCR4 D133N and R134A mutations provide quantitation for the most extreme known level of bias in CXCR4, through the direct and specific disruption of the interface between the receptor and the G protein while sparing the β-arrestin interaction. It is therefore noteworthy that the N-terminal Δ1-15 truncation mutant of CXCR4 shows nearly the same quantitative level of bias in our experiments. Again, this data challenges the notion that the receptor N-terminus is important exclusively for binding affinity; our data suggest that it not only affects signaling efficacy but also regulates signaling bias.

### Quantitative dissection of mutation effects benefits from amplification-free assays

A key distinction between this study and prior mutagenesis studies is the use of direct BRET-based interaction methods between CXCR4 and its effectors. As previously established (*43-45, 64-66*), for assays where measurements ultimately depend on amplified downstream second messengers, the magnitude of the measured signal is amplified as a result. The amplification of second messengers commonly measured in chemokine receptor mutagenesis studies is illustrated in fig. S1C for both IP3 & Ca^2+^, although the actual number of second messenger molecules is far greater than shown. Moreover, such measurements may reach their experimental maximum, either by saturation of the observation method (such as a fluorescent dye used to measure intracellular Ca^2+^), or by transient exhaustion of the intracellular second messenger stores or precursors themselves, long before full receptor occupancy and/or activation, so that differences between WT and mutant receptor activation are obscured.

To probe the utility of an amplification-based assay for quantitative dissection of mutation effects on CXCR4 signaling, we selected several CXCR4 mutations and truncations that showed pronounced defects in mini-G_αi_ BRET experiments, and tested them in Ca^2+^ mobilization experiments by constructing a CXCL12 concentration response curve for each (fig. S7A). Mutant surface expression was manually adjusted to closely match that of WT CXCR4 (fig. S7B). As expected, the Ca^2+^ mobilization experiments consistently yielded less pronounced mutation effects (fig. S7C-J), suggesting that these experiments are indeed subject to amplification when used to study chemokine receptors. This demonstrates that the quantitative nature of findings in our study was aided by the use of amplification-free, stoichiometric molecular association assays such as BRET-based β-arrestin-2 and G_αi_ recruitment above.

## Discussion

With CXCR4 being a prototypical CXC family receptor, CXCL12 being its only known endogenous agonist, and both of them playing pivotal roles in immune system homeostasis and in numerous cancers, CXCR4-CXCL12 is one of the most studied receptor-chemokine complexes biochemically. Extensive mutagenesis efforts directed at both the receptor and the chemokine have generated insight into the roles of a large number of residues; nevertheless, numerous uncertainties remain. First, there are many inconsistent reports on the relative contribution of individual residue interactions to the affinity or signaling capacity of the complex. This is likely because prior mutagenesis efforts largely relied on amplification-based second messenger assays conducted in a single-point rather than concentration-response format, an approach that often fails to detect all but the most dramatic mutant defects. In combination with mutation-induced variations in receptor expression and trafficking, this has prevented a systematic and quantitative dissection of the receptor-chemokine interface residues (Table 2). Second, even when data is consistent, the molecular basis for the impact of the mutations is unclear, as a structure of the complex has not been determined. Thus, the structural role and quantifiable functional impact of the various interaction epitopes, including the sulfotyrosinated extracellular N-terminus of the receptor, have remained cryptic.

In the present study, we addressed these methodological issues by using BRET-based methods for detecting stoichiometric association of CXCR4 with G_αi_ and β-arrestin-2, which are not subject to amplification; moreover, we designed the assays in a way that allowed exact quantitative interpretation despite variations in mutant expression levels. Additionally, our studies are guided and interpreted with a computationally constructed high-resolution 3D model of the CXCR4-CXCL12 complex. Built by a hybrid approach combining homology modeling, ab initio structural optimization, and experimentally-derived disulfide crosslinking restraints between chemokine and receptor (*31*), the model features the engagement of the full N-terminus of the receptor with CXCL12, and elucidates other key intermolecular interaction epitopes. Together with experimental results of this study, it provides the most comprehensive structural understanding of CXCR4-CXCL12 signaling thus far.

Key charge swap mutagenesis experiments established the accuracy of the overall model architecture. In these experiments, loss of function caused by mutation of a charged residue on one of the interacting proteins is rescued by a complementary mutation on the second protein. This method is inherently superior to traditional single-sided loss-of-function mutagenesis, because in addition to demonstrating that specific residues are important for the function of the complex, it is capable of showing whether they interact in the predicted pairwise manner. Using this method, we confirmed the predicted salt bridge between CXCR4 Asp^262^(6.58) and CXCL12 Arg^8^, and another one between CXCR4 Glu^277^(7.28) and CXCL12 Arg^12^. Not all interacting residue pairs in the CXCR4-CXCL12 complex are amenable to this strategy; for example, residues in CRS2 render CXCR4 completely inactive when mutated, and mutations of single residues in CRS1 do not produce signaling deficits that are substantial enough to rescue. Therefore, the data on the two interacting pairs Asp^262^(6.58)-Arg^8^ and Glu^277^(7.28)-Arg^12^ provide the strongest possible charge swap support for the model, particularly for the CRS1.5 and CRS2 regions. Concurrently, disulfide crosslinking results obtained in a separate study established the geometry of CRS1 and CRS0.5 (*31*).

The significance of the two intermolecular salt bridges validated by the charge swaps extends beyond simply supporting the predicted geometry of the CXCR4-CXCL12 complex. In fact, sequence alignments (*14, 67*) demonstrate a striking degree of conservation of the residues forming the first of the two salt bridges (CXCR4 Asp^262^(6.58) with CXCL12 Arg^8^) in the CXC receptors and chemokines, respectively, but not at all in other subfamilies (CC/CX_3_C/XC). This suggests that the identified salt bridge may be a universal CXC recognition anchor and a determinant of inter-subfamily selectivity. It is also remarkable that the virally encoded chemokine vMIP-II, a rare CC chemokine that binds to a CXC receptor, has an Arg at N-terminal position 7 that as demonstrated in the crystal structure (*14*), is engaged with CXCR4 Asp^262^(6.58) in a salt bridge, largely mimicking the one predicted and validated here. Therefore, the presence of an Arg in the proximal N-terminus and the resulting ability to form a salt bridge with the acidic residue in position 6.58 of the receptor may confer a cross-subfamily activity to CC chemokines. By contrast with Asp^262^(6.58) and Arg^8^, residues forming the second salt bridge are not conserved: CXCL12 Arg^12^ is unique among the CXC chemokines, while a Glu at position 7.28 is only found in CXCR3 and ACKR3 (the latter of which also binds CXCL12). Therefore, this second salt bridge likely contributes to the intra-CXC-subfamily selectivity for the CXCR4-CXCL12 complex.

Our experiments also established an important role of CXCR4 Arg^30^(NT), a key predicted feature of CRS1.5. Interestingly, a basic residue in the corresponding position is found in almost every CC (but not CXC or other subfamily) receptor, where it has been proposed to be important for coordination of the uniquely-shaped CC-motif backbone of the respective CC chemokines (*67*). It is thus possible that a basic amino-acid in this position confers CXCR4 with a CC-like recognition determinant, and facilitates its interaction with CC chemokine, vMIP-II.

Mutations in CXCR4 CRS2 often have large effects, and in some cases completely ablate CXCL12-mediated signaling. This is true not only for G protein assays, in agreement with earlier studies (*9, 30, 32, 47-51*), but also, as we demonstrated here, for β-arrestin recruitment. Our findings are consistent with the receptor binding pocket residues and the N-terminus of CXCL12 being the key drivers of signaling (*2, 6*). By contrast, single-point mutations in other epitopes, and particularly the receptor N-terminus, generally have less dramatic effects. Accordingly, the discovery of the impact of progressive N-terminal deletions of CXCR4 on the efficacy of G_αi_ and β-arrestin-2 recruitment was surprising. This discovery challenges the long-standing paradigm in which the receptor N-terminus serves as no more than a docking domain for the chemokine (*2, 6*). Even with truncations, none of the signaling defects were as severe as those produced by CRS2 mutations. However, the fact that they were observed at all suggests a new and important role for the N-terminus beyond the two-site hypothesis; specifically, that signaling amplitude depends on the extent to which the receptor N-terminus binds the chemokine. Notably, the efficacy variations resulting from N-terminal truncations were only detectable in an amplification-free, molecular-association-based assay, which emphasizes the importance of using adequate tools and readouts in mutant characterization.

There are two possible mechanisms for the observed effects of receptor N-terminal truncations. On the one hand, the N-terminus may be directly involved in the conformational change underlying receptor activation. Consistent with this hypothesis, there is a direct covalent (disulfide) bond between the receptor N-terminus and the ECL3 that effectively links the N-terminus to the two activation-related helices, TM6 and TM7. This N-terminus-to-ECL3 disulfide is highly conserved across chemokine receptors, suggesting that this mechanism may be common. Also, on the opposite side of the binding pocket, packing of CXCL12 against ECL2 and the ability of the receptor to close down around the chemokine, akin to other GPCRs and their ligands (*68–70*), may both depend on the receptor N-terminus locking down the chemokine globular core. This concords with our observation of the detrimental effects of charge reversal of the CXCR4 ECL2 tip, which produced results similar to truncating the N-terminus. Finally, the intramolecular association between the N-terminus and ECL2 of the receptor, stabilized by the bound chemokine, may play a role in establishing the correct signaling geometry.

As an alternative to the conformational mechanism, the receptor N-terminus may affect signaling efficacy indirectly by prolonging the residence time of CXCL12 on the receptor. This hypothesis is inspired by our recent study of ACKR3 (which also binds CXCL12), where impairment in β-arrestin-2 recruitment efficacy produced by N-terminal truncations in the receptor strongly correlated with an increase in chemokine dissociation rate (*71*). A mechanistic link between ligand dissociation rates and signaling efficacy has also been established for other GPCRs (*43, 72*); specifically, efficacy differences between agonists have been explained by receptor occupancy relative to the kinetics of the signaling process under study. In the case of ACKR3, one can argue that more rapid chemokine dissociation, or in other words a shorter receptor residence time, prevents the receptor from being phosphorylated by G protein receptor kinases and coupling to β-arrestin-2. Although we have not yet established similar kinetic off-rate assays for CXCR4, it is possible that the observed signaling differences between WT CXCR4 and the truncated receptor are at least partially due to changes in chemokine dissociation rate. Such a role of the N-terminus in slowing the off-rate would also provide a partial explanation for the increased affinity of CXCL12 for CXCR4 in the presence of G protein (*73*), similar to the generation of a G protein-mediated “closed conformation” of the β_2_AR which prevents the egress of ligands (*68*). The geometry of the interaction, wherein the N-terminus has an extended structure that wraps around the chemokine, would facilitate a scenario in which alterations like tyrosine sulfation along with allosteric effects from G protein coupling could readily modulate the receptor N-terminus-chemokine interaction, thereby influencing signaling responses.

The involvement of the distal N-terminus in signaling may also explain why a disulfide locked dimer of CXCL12 has reduced signaling efficacy in β-arrestin-2 recruitment (*41, 57*). As described earlier, the distal N-terminus of the receptor forms an anti-parallel β-sheet with the β_1_-strand of the chemokine in a manner mimicking the chemokine dimer interface; thus binding of the chemokine dimer would displace the receptor N-terminus (fig. S6), likely producing similar conformational and/or ligand off-rate differences (relative to the complex with monomeric CXCL12) as we suspect for the N-terminal truncations tested herein.

In summary, this study presents novel insights into the functional anatomy of the CXCR4-CXCL12 complex and the role of various epitopes in regulating the structure, ligand specificity and signaling responses of the receptor. Some of these findings, such as the conserved CXC-specific salt bridge and the importance of the N-terminus, are likely to be broadly applicable in the chemokine receptor family, and provide structural explanation for the previously observed effects of mutations and N-terminal mutations in other receptors (*74*).

## Materials and Methods

### DNA constructs and cloning

Human CXCR4 fused to renilla luciferase 3 (rluc3, otherwise known as rlucII) and GFP fused to human β-arrestin-2 (GFP-β-arrestin-2), both contained within a pcDNA vector, were kindly donated by Nicolaus Heveker, Université de Montréal, Montreal, Quebec, Canada. An N-terminal HA tag was added to the CXCR4-rluc3 vector, followed by the production of our mutant library. All mutations and truncations (as well as the N-terminal HA tag) were introduced into the CXCR4-coding region of the CXCR4-rluc3 vector using the QuikChange site-directed mutagenesis method (Stratagene). The plasmid used to express renilla GFP (rGFP) fused to mini-G_αsi_ for the G_αi_ association BRET assay was a kind gift from Nevin Lambert, Augusta University, Augusta, Georgia, USA.

### BRET-based β-arrestin-2 and mini-G_αi_ association assays

β-arrestin-2 recruitment was measured with the bioluminescence resonance energy transfer 2 (BRET^2^) assay (*75*). Four days prior to each assay, HEK293T cells, cultured in Dulbecco’s Modified Eagle Media (DMEM) + 10% fetal bovine serum (FBS), were passaged and plated at 4.25×10^5^ cells per well in 6-well tissue culture plates.

For β-arrestin-2 association experiments, the cells were transfected two days later, with 0.1 µg DNA/well HA-CXCR4-rluc3 (WT and mutants, with 0.075 ug DNA used for the highly expressing Δ1-10 truncation) and 2.4 µg DNA/well GFP-β-arrestin-2. Transfections were carried out according to the manufacturer’s recommended protocol using TransIT-LT1 transfection reagent (MirusBio, Madison, Wisconsin, USA). For G_αi_ association experiments, the procedure was identical except that rGFP-mini-G_αsi_ was used in place of GFP-β-arrestin-2. On the day of the assay, cells were washed while still adherent with PBS, then re-suspended through manual pipetting in PBS + 0.1% D-glucose (BRET buffer) and diluted to obtain a final concentration of 1.5×10^6^ cells/mL of suspension. Ninety µL of cell suspension was then dispensed into each well of a white, clear bottom, tissue culture treated 96-well plate (Corning). The plate was placed into a 37°C CO_2_ conditioned incubator for 30 min before GFP-β-arrestin-2 or rGFP-mini-G_αsi_ fluorescence levels were measured with a SpectraMax M5 fluorescence plate reader (Molecular Devices, San Jose, California, USA). For β-arrestin-2 experiments, 10 µL of BRET buffer containing 10 times the final intended concentration of CXCL12 (WT or mutant) was then added to each well, and the plate was placed in a VictorX Light multilabel plate reader (PerkinElmer Life Sciences, Waltham, Massachusetts) warmed to 37°C for 10 min. Coelenterazine-200A (a.k.a. Deep Blue C) was then added to each well in order to obtain a final concentration of 5 µM immediately before repeatedly measuring the luminescence at both 410 and 515 nm in the VictorX Light luminometer. In the case of G_αi_ association experiments, the coelenterazine-200A was added immediately *before* CXCL12, and the luminescence was measured repeatedly immediately after adding CXCL12. BRET ratios (515 nm luminescence/410 nm luminescence) (from 20 min after CXCL12 addiction for β-arrestin-2 experiments, 1 min for G_αi_) were then calculated using MS Excel, and initial 4-parameter agonist concentration response curve fitting was carried out using GraphPad PRISM in order to enable normalization to WT E_max_ within each experiment.

Vector amounts used in the BRET experiment transfections, 0.1 and 2.4 µg for HA-CXCR4-rluc3 and GFP-β-arrestin-2/rGFP-mini-G_αsi_ respectively, were selected to meet three criteria. First, the expression of GFP-effector relative to CXCR4-rluc3 was high enough to ensure the potential for saturation of CXCR4-rluc3. Experiments were designed in this way to avoid misinterpretation of apparent efficacy changes resulting purely from expression-based deviations from potential “BRET_max_”, or the upper limit of the hyperbolic donor:acceptor titration relationship observed for interacting BRET-donor and acceptor-fused pairs (*75*). The second criterion met was that a sufficient level of rluc3 must be expressed in order to yield an analyzable signal for both wavelengths of luminescence measured in the BRET assay, and the third was simply that the maximum DNA recommended DNA for the scale used (2.5 µg/well) was transfected into the cells.

The orientation of our BRET assays, with receptor linked to the energy-donating luminescent enzyme and saturated by a signaling effector fused to the accepting fluorescent protein, provides the advantage of rendering the experiments insensitive to moderate variations in receptor expression levels. This is due to the ratiometric BRET signal representing the proportion of donor in close proximity to acceptor, rather than the absolute quantity of engaged complexes. As long as there is ample GFP-effector available (enabled by saturation), the proportional BRET signal should represent the per molecule average receptor-effector engagement or complex rearrangement within a sample.

To confirm this, we titrated WT CXCR4-rluc3 in both the β-arrestin-2 and mini-G_αsi_ BRET experiments, and indeed no significant changes in signaling parameters were observed (fig. S2). The range of total WT CXCR4-rluc3 expression that produced functionally equivalent activation curves encompasses the expression seen for all mutants tested herein except for CXCR4(Δ1-10) (fig. S3A&D), which was expressed at higher than WT levels. It should be noted that the proportional surface expression is altered to varying degrees for a number of mutants (fig. S3C&F), and in severe cases there may indeed be an effect on BRET measurements, as a smaller proportion of the donor-fused receptor being available to CXCL12 at baseline would be expected to produce a lower proportional saturation of donor by acceptor upon stimulation (*56*). We therefore note every such case in the results (for β-arrestin-2 BRET: Δ1-19, D133N, R134A; for mini-G_αsi_: Δ1-15, Δ1-25, D133N, R134A), and we interpret these particular data with caution to the degree warranted by the surface expression deficit.

### Statistical comparison of WT and mutant signaling parameters

In order to compare WT and mutant signaling assay results, as well as the results of different combinations of CXCL12 and CXCR4 mutants, results on each day were normalized to 100% WT efficacy, and mean values from independent experiments (each performed in duplicate) for each CXCL12 concentration were plotted together. Statistical comparisons between WT and mutant CXCR4 concentration response parameters were carried out on the same combined dataset in Graphpad Prism version 5.0b for Mac OS X (GraphPad Software, San Diego, California, USA, www.graphpad.com) using the Akaike’s informative criteria (AICc) test, the results of which are contained in Table 3. It should be noted that the statistical test used to establish differences between WT and mutant CXCR4 signaling parameters was in some cases inadequate for assigning significance simply due to near complete elimination of receptor activation upon mutation. For example, the efficacy of CXCR4(Δ1-25) is not significantly different in efficacy from WT CXCR4 in mini-G_αi_ association experiments according to the AICc test used, which is clearly incorrect and arises from the severely impaired efficacy seen for Δ1-25 and the resultant failure to reach saturation.

### Ca2+ mobilization G protein signaling assay

Ca^2+^ mobilization experiments were carried using the FLIPR4 calcium assay dye kit (Molecular Devices). As detailed previously (*32*), we use a modified CHO-K1 cell line for these experiments that stably expresses the promiscuously coupling G_α15_. These cells were cultured in 1:1 DMEM/F12 nutrient mixture supplemented with 10% FBS and 700 µg/mL active G418 mammalian antibiotic. Three days prior to each assay, the cells were passaged and plated at 2×10^5^ cells per dish into 10 cm diameter tissue culture dishes, in 1:1 DMEM/F12 nutrient mixture supplemented with 10% FBS and further supplemented with 0.25% DMSO to aid in transfection efficiency (*76*).

The next day, the media was removed and replaced with 1:1 DMEM/F12 nutrient mixture supplemented with 10% FBS immediately prior to transfection with Trans-IT CHO transfection kit, which was used according to the manufacturer’s recommended protocol except that the µL reagent:µg DNA ratio was adjusted to 4:1. For the current study, 24 ug of either WT or mutant CXCR4-rluc3 DNA were transfected into the cells in each dish. We used the same WT and mutant CXCR4-rluc3 constructs as were used in BRET experiments, as we discovered that CXCR4-rluc3 retained activation of Ca^2+^ mobilization in the CHO-K1-G_α15_ cell line and seemed to improve the data quality (fig. S7A). In the case of Ca^2+^ mobilization experiments, expression levels do affect signaling measurements substantially, so WT CXCR4 expression levels were adjusted to match those of mutants as closely as possible using flow cytometry-based monitoring of anti-HA-PE or anti-HA-APC binding to the N-terminal HA (fig. S7B).

The following day, the cells were washed with PBS before re-suspension using PBS + 5 mM EDTA, then centrifuged and re-suspended in 1:1 DMEM/F12 nutrient mixture supplemented with 10% FBS before re-plating at 90,000 cells/well in black, clear bottom, poly-D-lysine coated 96-well plates (Corning).

Finally, on day four, the media was carefully removed from the adherent cells before 200 µL of a 1:1 mixture of HBSS + 20 mM HEPES + 0.1% BSA (Ca^2+^ flux buffer) and FLIPR4 dye was added to each well. After a 75 min incubation at 37°C, the assay was carried out in a FlexStation-3 multi-mode plate reader (Molecular Devices) using the instruments automated injection function to inject CXCL12 at 10 times the final indicated concentrations (in Ca^2+^ flux buffer) while reading fluorescence (emission 525 nm; excitation 485 nm) repeatedly (with 1.52 second intervals) over the course of 150 seconds. Reduced peak fluorescence values were calculated by subtracting baseline fluorescence from peak values. Reduced peak fluorescence 4-parameter CXCL12 concentration response curve fitting was carried out using GraphPad PRISM.

### Flow cytometry-based surface expression testing

Cell surface expression of WT and mutant versions of CXCR4-rluc3 was monitored by flow cytometry as described previously (*32*). Briefly, cells were resuspended in PBS + 5 mM EDTA, centrifuged, and re-suspended in PBS + 0.5% PBS (FACS buffer) to a final concentration of 0.1-1×10^6^ cells/mL of buffer. For anti-HA staining, fluorophore-conjugated anti-HA antibody (either anti-HA-APC, catalogue number 130-098-404, or anti-HA-PE, catalogue number 130-092-257, both Miltenyi Biotec, Bergisch Gladbach, Germany) was added to obtain an 11X dilution, and cells were stained on ice in the dark for 10 min, according to the manufacturer’s recommended procedure. For anti-CXCR4 staining, fluorophore-conjugated anti-CXCR4 antibody, either 12G5 anti-CXCR4-APC (catalogue number 560936, BD Biosciences, La Jolla, California, USA) for N-terminal truncations and CRS1 mutations or 1D9 anti-CXCR4-PE (catalogue number 551510, BD Biosciences) for all other mutations was added to obtain a 50X dilution, and cells were stained on ice in the dark for 45 min, as recommended by the manufacturer. Cells were then washed three times with FACS buffer and then fixed with a final concentration of 0.8% PFA. Flow cytometric analysis of the antibody-stained and fixed cells was carried out using a GUAVA benchtop flow cytometer (EMD Millipore, Burlington, Massachusetts, USA). Flow cytometry data analysis was performed using FlowJo version 10 (FlowJo LLC, Ashland, Oregon, USA), and geometric mean fluorescence intensity (GMFI) data were normalized to that of WT CXCR4 after subtraction of the low GMFI obtained for pcDNA-transfected control cells stained with the same antibody.

### Bias calculations

Bias was calculated according to the equiactive comparison method for bias calculation (*45*), which requires only two pairs of EC_50_ and E_max_ parameters from assays of two different GPCR-initiated signaling pathways. While this equation is typically used to assess bias between different agonists, it was perfectly suited to our case of seemingly differential mutational effects on β-arrestin-2 and G_αi_ association. The adapted equation is:

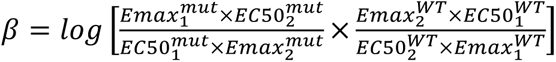

where 1 and 2 correspond to pathway 1 and 2, designated arbitrarily. As G protein signaling is usually considered the primary function of GPCRs, we designated G_αi_ association as pathway 1 and β-arrestin-2 association as pathway 2, so that negative results indicate a bias towards β-arrestin-2 association. The signaling parameters were applied to the above adapted equiactive bias equation, and the error of the parameters combined, in MS Excel.

## Acknowledgments

We gratefully acknowledge N. Lambert (Augusta University), M. Bouvier (Université de Montréal), and M. Gustavsson (UC San Diego) for BRET reagents used in the proposed studies. We thank J.J. Ziarek (Indiana University) and B.F. Volkman (Medical College of Wisconsin) for providing the coordinates of their previously published CXCR4-CXCL12 model.

## Author contributions

T.M.H. and I.K. conceived the study and provided overall supervision of the project. B.S.S., T.M.H., and I.K. designed experiments. B.S.S. designed and cloned receptor and chemokine constructs, prepared purified chemokine variants, performed all experiments, and analyzed the data. B.S.S., T.N., T.M.H., and I.K. interpreted the data in the context of the model. B.S.S, T.N., T.M.H. and I.K. wrote the paper, with all authors providing input and approving the final version.

## Data and materials availability

The coordinates of the CXCR4-CXCL12 complex utilized in the present study are available upon request.

## Supplementary Materials

**Fig. S1.**
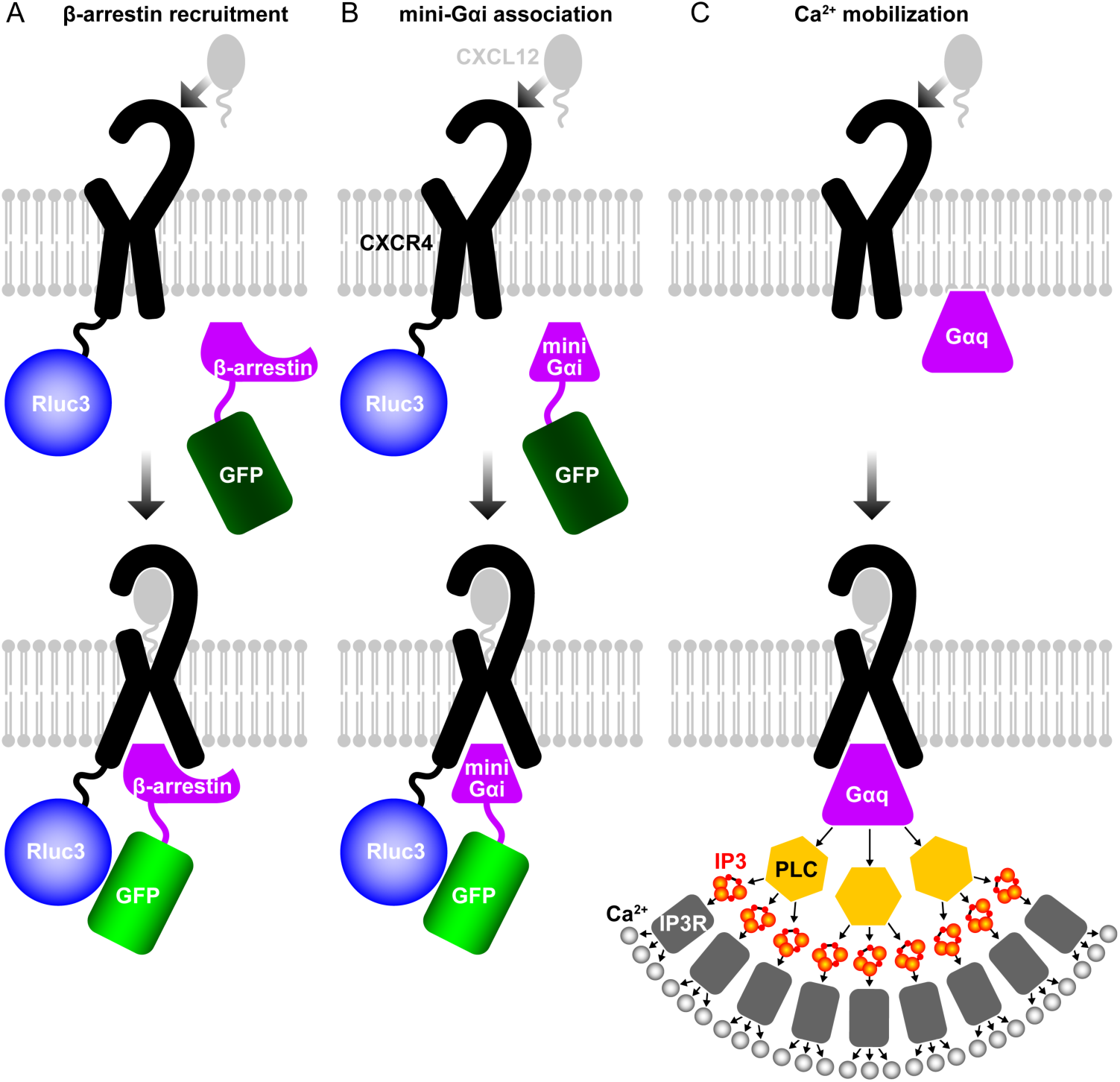
Schematics of assays used for characterization of CXCR4 mutants in this study. (**A**) The BRET-based β-arrestin-2 recruitment assay employs the receptor that is C-terminally tagged with engineered Renilla luciferase 3 (rluc3, also known as rlucII), and a β-arrestin-2 protein tagged with GFP. Upon ligand stimulation, β-arrestin-2 is recruited to the activated receptor, which brings the two tags in close proximity and results in energy transfer from rluc3 (donor) to GFP (acceptor), detectable as the ratio of emission at 515 nm to the emission at 410 nm. This assay detects one-to-one association between the receptor and β-arrestin-2, with no signal amplification. **(B)** The BRET-based mini-G_αi_ association assay works by the same principle as in (A), except that the acceptor is fused to an engineered G_αi_ protein, mini-G_αi_. Similarly to (A), this assay detects one-to-one association between the receptor and mini-Gαi, with no signal amplification. (**C**) The intracellular Ca^2+^ mobilization assay relies on a sequence of intracellular events that convert the activation of the receptor to the production of the second messenger IP3 which in turn stimulates the release of the second messenger Ca^2+^ from intracellular stores. The activation of PLC by G_αq_, the production of IP3 by the activated PLC, and the release of Ca^2+^ upon IP3 binding to the intracellular IP3 receptor (IP3R) involve signal amplification.

**Fig. S2.**
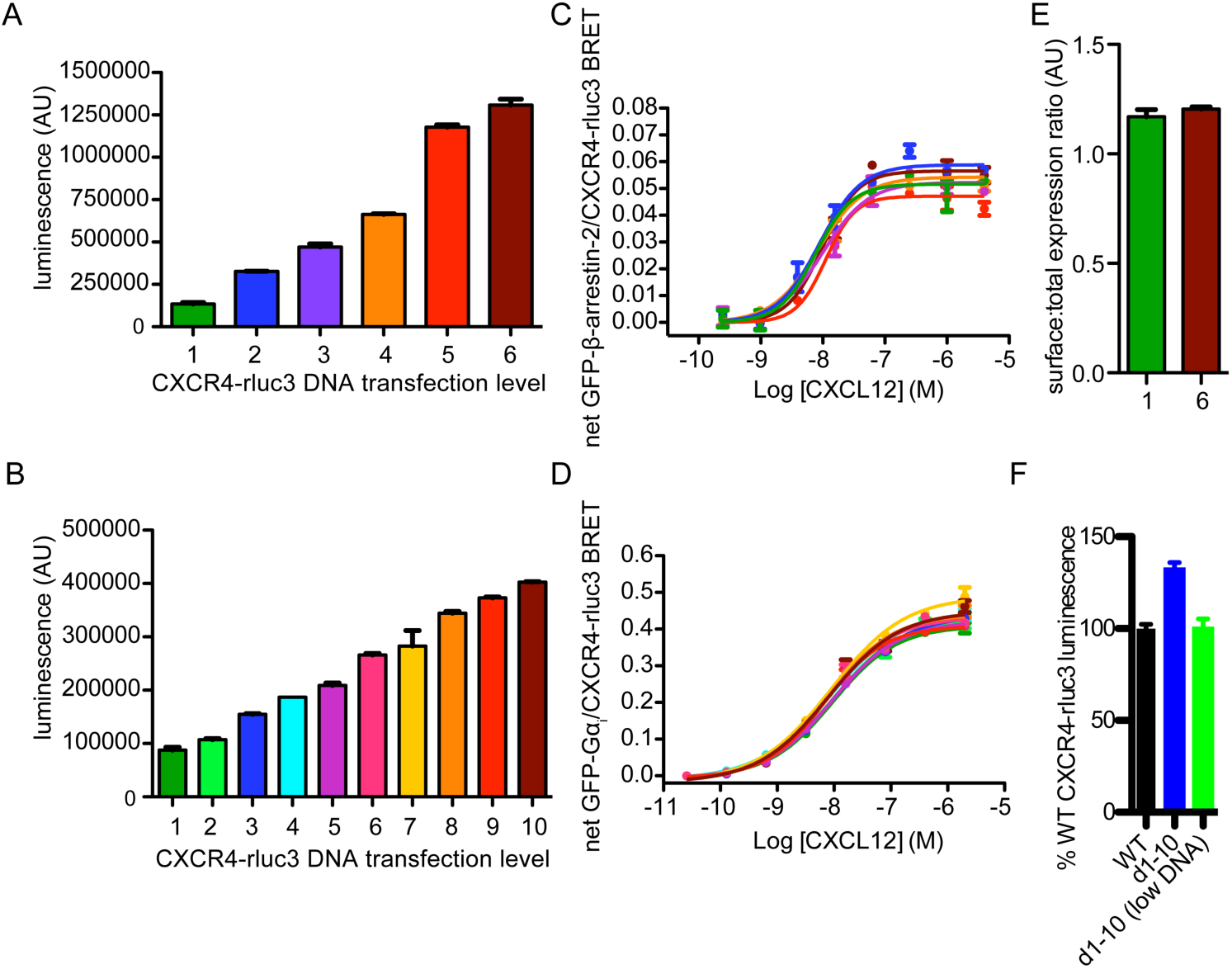
CXCR4 yields identical CXCL12-mediated β-arrestin-2 and G_ɑi_ signaling parameters in BRET experiments across a wide range of expression levels. (**A**-**B**) Luminescence of HEK293T cells 48 hours after transfection with various amounts of CXCR4-rluc3 DNA, along with an equal high level of (**A**) GFP-β-arrestin-2 or (**B**) rGFP-mini-G_ɑi_, extending above and below the amount of WT or mutant CXCR4-rluc3 DNA used in the various other BRET experiments herein. (**C**) Net CXCL12-mediated BRET ratio data for the samples in (**A**) after stimulation with varying CXCL12 concentrations for 20 minutes. (**D**) Net CXCL12-mediated BRET ratio data for the samples in (**B**). In the case of both experiments, no significant differences between EC_50_ and E_max_ parameters were found between the different samples by the Akaike’s informative criteria (AICc) test. (**E**) Surface:total CXCR4 expression ratio of the highest and lowest CXCR4-rluc3-expressing cells from (**A**). Surface expression of CXCR4 was determined by flow cytometry-based measurement of anti-CXCR4(12G5)-PE binding. (**F**) Luminescence of cells expressing adjusted (lowered) and pre-adjustment levels of Δ1-10 CXCR4-rluc3, normalized to that of cells expressing WT CXCR4-rluc3, tested in the same experiment. In all cases, data from one independent experiment representative of three are shown.

**Fig. S3.**
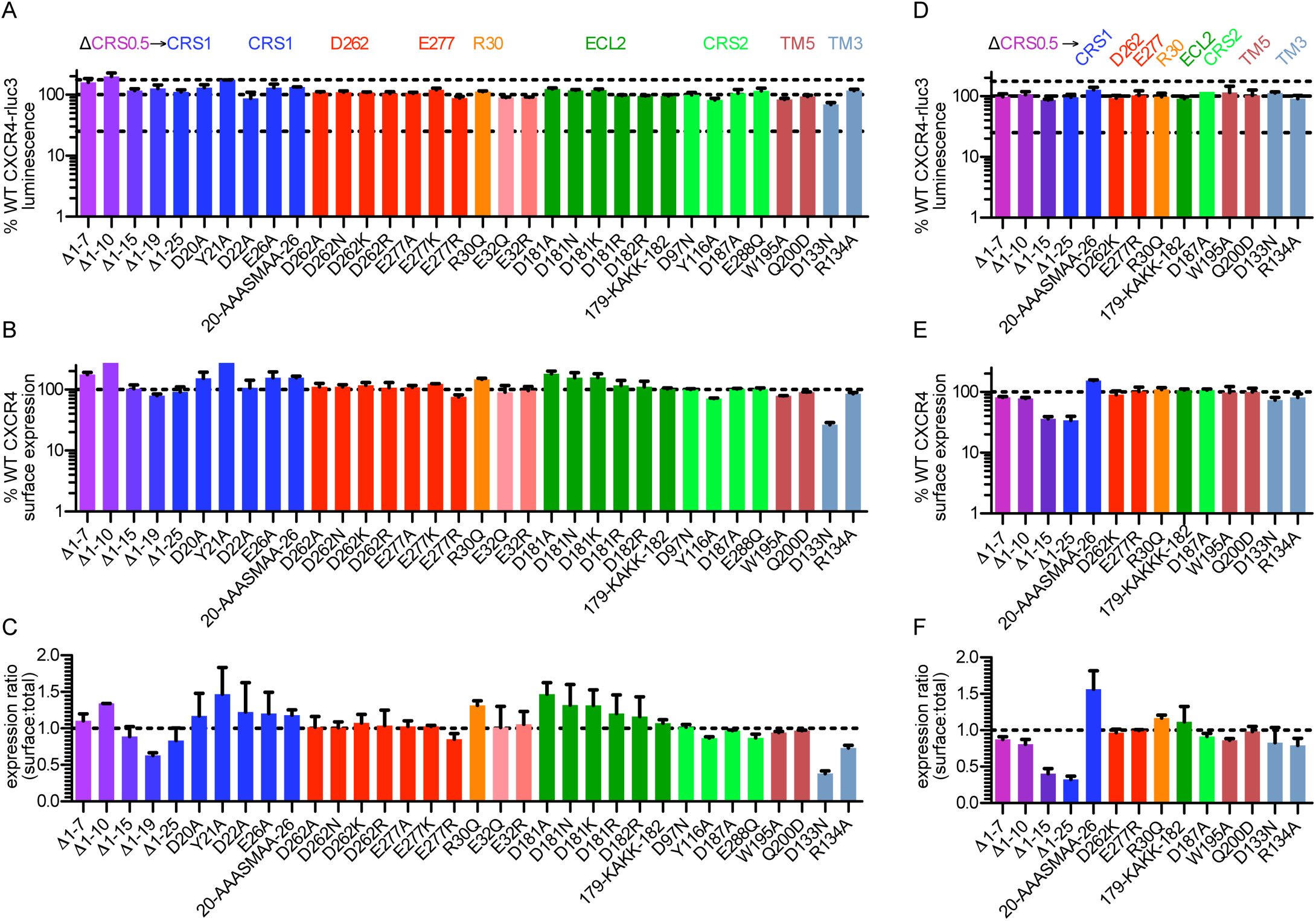
Total and surface expression of mutants of residues whose importance for signaling was suggested by BRET experiments. (A) Total expression of CXCR4 mutants when co-expressed with β-arrestin-2, as determined by the rluc3 luminescence of the expressing cells. (B) Surface expression of CXCR4 mutants when co-expressed with β-arrestin-2, as determined by flow cytometry based detection of anti-CXCR4-PE or anti-CXCR4-APC binding. For N-terminal perturbations, the 12G5 (ECL2 epitope) anti-CXCR4 antibody was used, whereas for TM domain mutations, we used the 1D9 (N-terminal epitope) antibody. (C) The surface:total expression ratio, or the ratio of the data in (B) to those in (A). (D-F) Same as in (A-C) respectively, except mutants were tested when co-expressed with rGFP-mini-G_ɑi_. All data are the mean of two independent experiments, each performed in duplicate with data normalized to WT CXCR4 results from the same experiment.

**Fig. S4.**
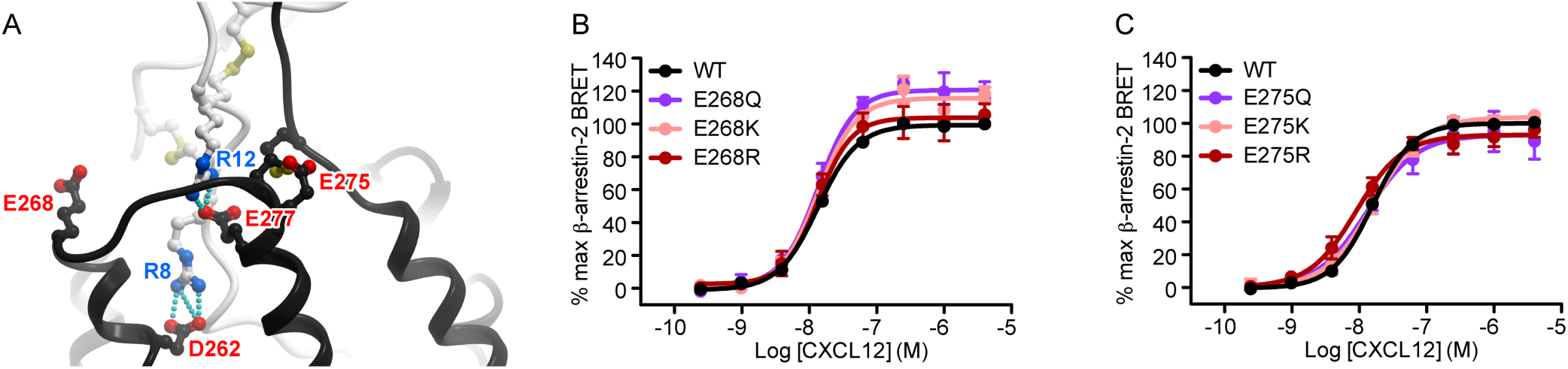
CXCR4 residues Glu^268^(ECL3) and Glu^275^(7.26) are not critical to CXCL12-mediated CXCR4 activation. (**A**) The location of two additional CXCR4 ECL3 acidic residues, Glu^268^(ECL3) and Glu^275^(7.26), relative to the predicted and experimentally-validated salt bridges between CXCR4 Asp^262^(6.58) or Glu^277^(7.28) and CXCL12 R^8^ or R^12^, respectively. (**B**-**C**) β-arrestin-2 association BRET ratio data for (**B**) a series of CXCR4 Glu^268^(ECL3) mutants (E268Q/K/R) or (**B**) a series of CXCR4 Glu^275^(7.26) mutants (E275Q/K/R) after stimulation with varying CXCL12 concentrations for 20 minutes. Data represent the mean values (with error bars indicating SEM) from at least three independent experiments, each performed in duplicate with data normalized to the E_max_ of WT CXCR4 tested in the same experiments.

**Fig. S5.**
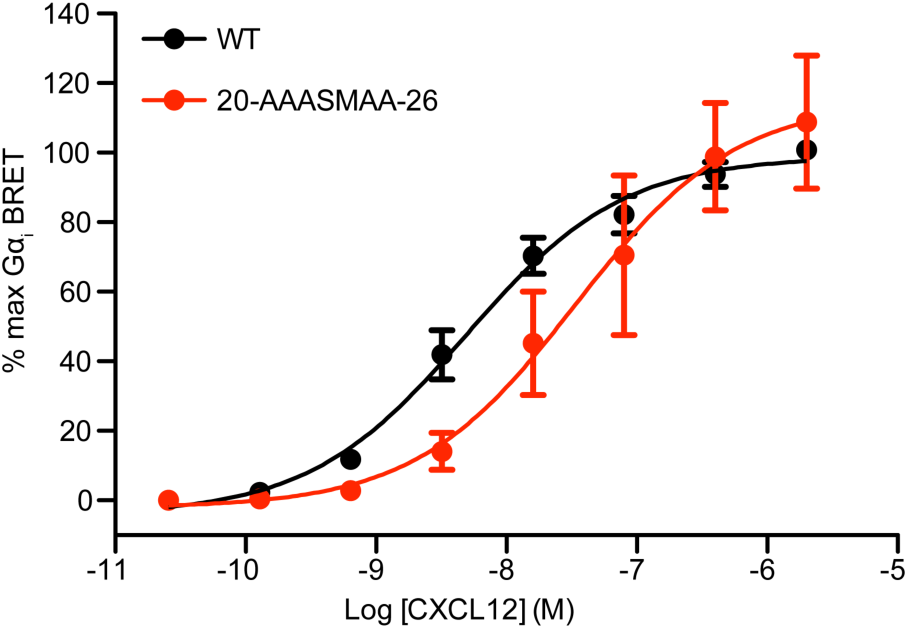
Negatively charged residues in CRS1 are important to the potency of CXCL12-mediated CXCR4-G protein engagement. Mini-G_ɑi_ association BRET ratio data is shown for CXCR4(20-AAASMAA-26) after stimulation with varying CXCL12 concentrations for 1 minute. Data represent the mean values (with error bars indicating SEM) from at least three independent experiments, each performed in duplicate with data normalized to the E_max_ of WT CXCR4 tested in the same experiments.

**Fig. S6.**
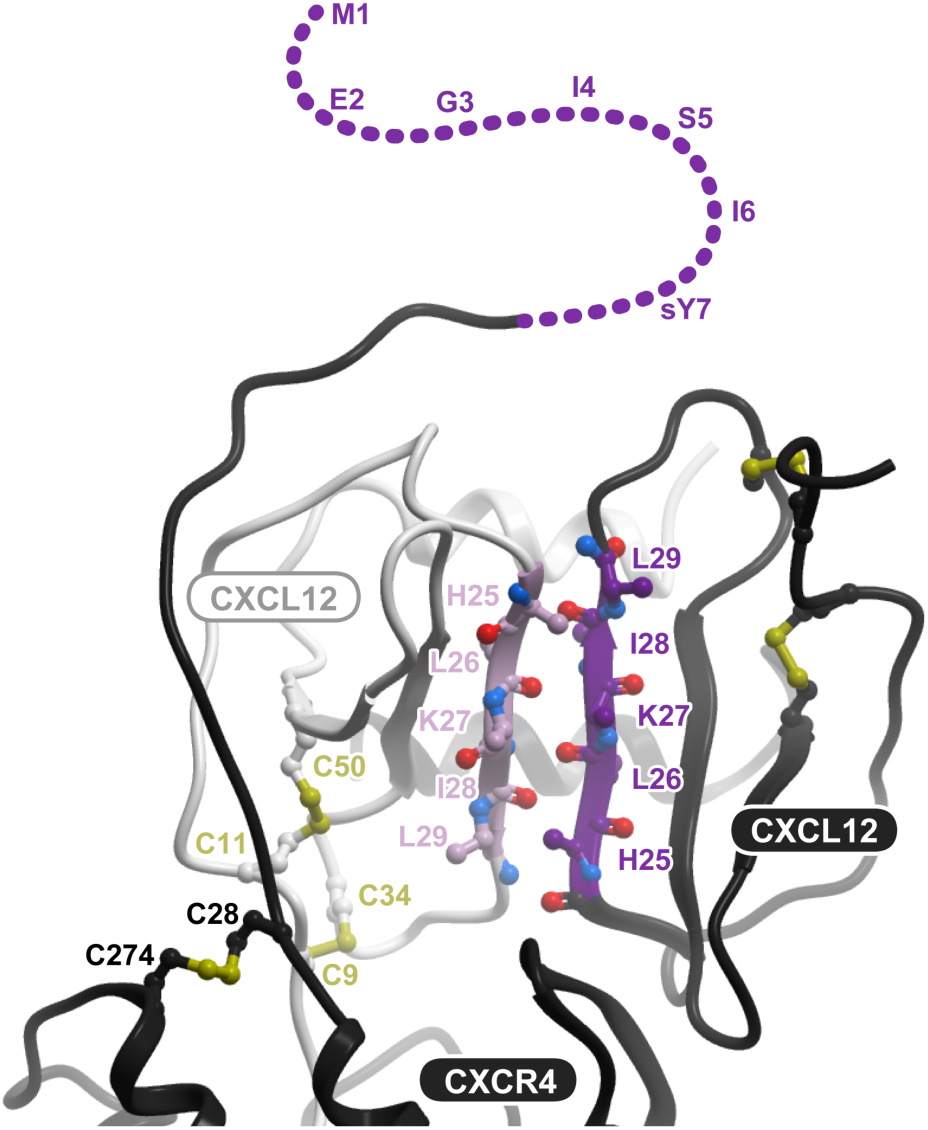
Predicted consequence of dimeric CXCL12 binding to CXCR4 in the context of our model. Binding of CXCR4 to dimeric CXCL12 is sterically feasible and preserves CRS1, CRS1.5, and CRS2 interactions similar to monomeric CXCL12; however, it is mutually exclusive with the proposed CRS0.5 interaction (compare to Fig. 1B).

**Fig. S7.**
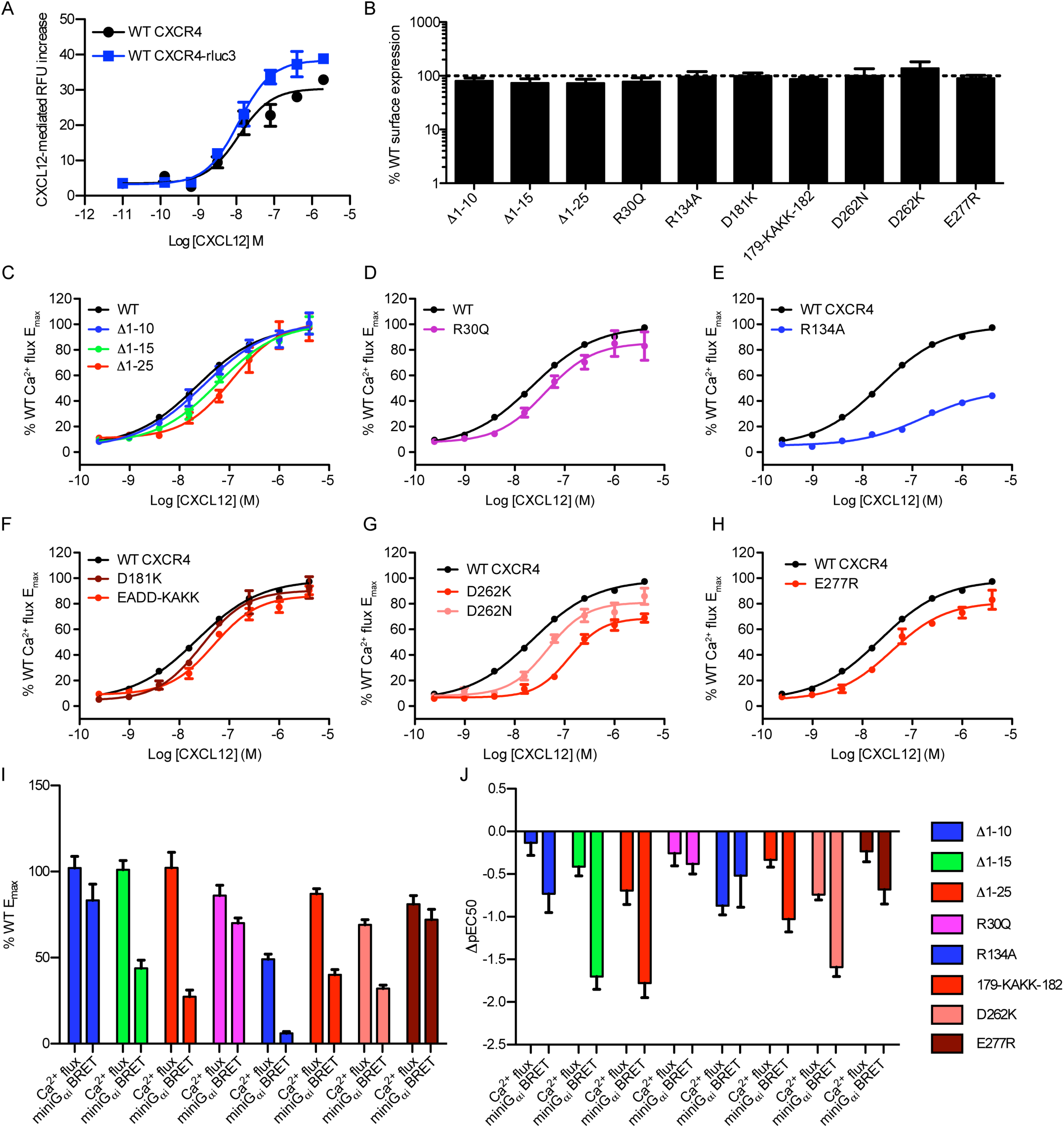
Signal amplification in Ca^2+^ mobilization experiments obscures mutation-induced defects. (**A**) Ca^2+^ mobilization (Ca^2+^ flux) in CHO-K1-G_α15_ cells expressing WT HA-CXCR4 with no C-terminal fusion or WT HA-CXCR4-rluc3 upon stimulation with varying CXCL12 concentrations. Data with the baseline RFU value subtracted is shown from one independent experiment representative of two. (**B**) The expression of mutants tested in Ca^2+^ flux, as determined by flow cytometry based detection of anti-HA-PE or anti-HA-APC binding, monitored in the same cells as measured for Ca^2+^ flux and normalized to the WT CXCR4 expression level from within the same experiment. For all mutant surface expression data, N ≥ 3 and error bars represent SEM. (**C**-**H**) Ca^2+^ mobilization (Ca^2+^ flux) in CHO-K1-G_α15_ cells expressing (**B**) CXCR4(Δ1-10), CXCR4(Δ1-15), or CXCR4(Δ1-25), (**C**) CXCR4(R30Q), (**D**) CXCR4(R134A), (**E**) CXCR4(D181K) or CXCR4(179-EADD-182), (**F**) CXCR4(D262K) or CXCR4(D262N), or (**G**) CXCR4(E277R) upon stimulation with varying CXCL12 concentrations. Ca^2+^ flux data represent the mean values (with error bars indicating SEM) from at least three independent experiments, each performed in triplicate with data normalized to the E_max_ of WT CXCR4 tested in the same experiments. (**I**-**J**) WT-normalized E_max_ **(I)** and ΔpEC_50_ **(J)** parameters from Ca^2+^ mobilization experiments in (**C**-**H**) next to those from the mini-G_αi_ BRET experiments used to test the same mutations.

